# Time-resolved phylogenomics analysis reveals patterns in biosphere nutrient limitation through Earth history

**DOI:** 10.1101/2025.11.13.688358

**Authors:** Zhanghan Ni, Troy Osborn, Juntao Zhong, Adrián González, William Puzella, Aya Klos, Bryan Le, Alec Leonetti, Joanne Boden, Eva Stueeken, Rika Anderson

## Abstract

The co-evolution of life and Earth has profoundly transformed global biogeochemical cycles over the past 3.5 billion years. These cycles, in turn, have dictated the availability of essential nutrients like phosphorus, nitrogen, and iron, thereby affecting primary productivity and the scale of the Earth’s biosphere. Despite the critical role of nutrient limitation in shaping the size and scope of the biosphere, significant uncertainties persist about which nutrients were globally limiting at various points in Earth history. Here, we use a phylogenomic approach to trace the origin and spread of genes associated with nutrient limitation over time. We show that genes associated with phosphorus limitation emerged relatively early in life’s history, whereas genes associated with nitrogen limitation emerged later, closer to the Great Oxidation Event. In terms of iron limitation, we present novel evidence that siderophores, compounds that facilitate iron uptake, may have arisen as early as the Archean. Overall, our results have important implications for understanding how the geosphere has influenced the scale and extent of life on Earth for the past 4 billion years.

## Introduction

For nearly 4 billion years, the co-evolution of life and Earth has driven profound transformations in planetary biogeochemistry (*1*). This has led to dynamic fluctuations in the magnitude and rate of primary production on a planetary scale over time (*2–4*). The modern biosphere, encompassing an estimated 700 billion tons of carbon (*5*), exhibits rapid carbon cycling through the interconnected oceanic, terrestrial, and atmospheric reservoirs, with an estimated turnover time of less than 8 years (*6*, *7*). However, great uncertainty remains regarding how much life the combined geobiosphere was able to sustain at different periods throughout Earth history (*4*).

The availability of nutrients and energy (i.e. electron donors and acceptors) exerts a primary control on the overall productivity of the biosphere. Essential elements, including phosphorus, nitrogen, iron, and sulfur, are critical for fundamental biological processes like biomass production and/or energy metabolism. However, their availability has exhibited significant fluctuations throughout Earth’s history, profoundly impacting the abundance and distribution of life (*2*, *8–11*). Notably, phosphorus cycling is commonly proposed to impose upper limits on the size and impact of the biosphere on a global scale (*12*, *13*), but nitrogen and iron are also commonly limiting nutrients in modern ecosystems (*11*). The availability of ferrous iron may have limited the size of the early Archean biosphere (from approximately 4.0-2.5 billion years ago (Ga)), when primary producers were perhaps dominated in part by photoferrotrophs (*14*). In the modern ocean, productivity tends to be limited by nitrogen in low-latitude equatorial regions, whereas iron is more limiting where subsurface nutrient availability is higher, such as in the Southern Ocean (*15*). The question of which nutrients would have been limiting on a global scale throughout the Archean (from 4.0 to 2.5 Ga) and Proterozoic (from 2.2 to 0.54 Ga) remains open, with unresolved implications for global primary productivity, the biological carbon pump, and carbon burial (*2*, *10*, *16*).

In response to nutrient and substrate limitation, organisms can scavenge elements by expressing specific proteins that increase the rate of uptake from the surrounding medium (*17*). For example, lineages of the marine cyanobacterium *Prochlorococcus* living in phosphate-limited regions encode more phosphate transporters in their genomes than *Prochlorococcus* lineages inhabiting phosphorus-replete regions (*18*, *19*). More generally, Lockwood et al. found that plankton in phosphate-depleted regions encode more genes to scavenge P and C from phosphonates (*20*). Microbes can also alter their elemental stoichiometry to adapt their intrinsic nutrient requirements according to environmental availability (*10*). For example, phospholipids can be replaced with sulfolipids in microbial cellular membranes when phosphorus is scarce (*21*). Similarly, lineages of the cyanobacterium *Synechococcus* encode fewer iron-containing nitrate/nitrite reductase proteins in iron-deficient regions (*22*). These and other studies have demonstrated that the environment shapes microbial genomes such that the genomic content of organisms often reflects the nutrient limitations they have encountered (*19*, *22*, *23*).

Both geochemical and modeling approaches have been employed to better understand the history of nutrient and energy substrate limitation on Earth. Biogeochemical models can be used to reconstruct element cycles over long timescales (e.g. (*12*, *24*, *25*), informed by geochemical analyses of the rock record, laboratory experiments and insights from modern analogues (*12*). However, the rock record is often incomplete or ambiguous, making interpretation challenging. For example, estimates of phosphate concentrations in Archean seawater diverge over five orders of magnitude (reviewed in (*26*), hindering our ability to accurately estimate phosphorus availability for early life. A new approach is needed to overcome these limitations.

Microbial genomes offer a valuable complementary perspective. They act as a historical record of microbial adaptation over time, reflecting the genomic strategies that organisms have developed to adapt to nutrient-poor or nutrient-replete environments. Recent studies have identified patterns of nutrient limitation in global oceans based on gene presence and absence in microbial genomes (*19*). Here, we leverage large datasets generated by modern metagenomics studies to identify genes that effectively act as “sensors” for nutrient limitation on the modern Earth. We then reconstruct the evolutionary history of those genes to gain insights into which elements have limited primary productivity in deep marine, shallow marine, and terrestrial habitats throughout Earth history. In doing so, we place new constraints on how the biosphere has responded to fundamental changes in planetary biogeochemistry over time.

## Results and Discussion

### Identification of “sensor” genes for nutrient limitation

To identify genes that can effectively act as indicators of nutrient limitation, we used a two-pronged strategy: one grounded in the literature, and one grounded in a statistical analysis. For the literature-focused approach, a thorough review was conducted to create a manually curated list of genes associated with scavenging nitrogen, phosphorus, and iron. Our final list contained a total of 10 genes (clustered into Clusters of Orthologous Groups, or COGs (*27*)) associated with iron scavenging and limitation for both ferrous and ferric iron (7 associated with Fe^3+^, 3 associated with Fe^2+^), 15 COGs associated with phosphorus limitation, and 4 COGs associated with nitrogen limitation (**Supplementary Table 1**) based on previously published results. While many genes have been identified as being associated with nutrient limitation, we were deliberately conservative in constructing our final list of COGs. COGs identified through our initial literature review were eliminated if they failed to pass several criteria: first, if the COG included genes with functions that were not strictly associated with nutrient limitation; and secondly, COGs were eliminated if the genes were associated with scavenging nutrients other than the element in question. For example, MntH is an iron transporter that is also associated with uptake of manganese; our goal was to include genes that can be used as an indicator of specific nutrients.

The final list of genes associated with iron limitation includes key components of iron uptake and transport, such as outer membrane receptors for ferric siderophores (e.g. EntCDEF, FepA, TonB, FecA, and FhuE) and transport proteins like FeoA. The gene list for phosphate limitation includes important phosphorus transporters such as alkaline phosphatases and the Pho regulon (e.g.. PhoABDX) and ABC-type phosphate (i.e. PstS) and phosphonate transporters (e.g. PhnCEGHILMNJ). The list for nitrogen limitation includes genes related to nitrogen uptake and assimilation, including nitrate/nitrite transporters (e.g. FocA), cyanate degradation (i.e. CynS), and enzymes such as urease (e.g. UreE), and we also elected to include nitrogenase (nifD). For more details, see **Methods** and **Supplementary Figure 1**.

However, identifying genes from the literature is subject to biases and limitations; we are restricted to genes that other studies have chosen to focus on and are likely limited to specific taxa. Furthermore, some genes may have additional functions that are as of yet unknown. Thus, identifying genes based on the literature is inherently limited in scope. To use a more agnostic approach to identify genes that can act as indicators of nutrient stress, we used a statistical model to identify genes whose abundances are negatively correlated with concentrations of phosphate, nitrate/nitrite, and iron in modern oceans (see **Methods and Supplementary Figure 1**). To do this, we used data from the Tara Oceans dataset (*28*, *29*), which includes the relative abundances of genes within microbial communities (classified as COGs, or Clusters of Orthologous Groups (*27*)) alongside the concentrations of nutrients like phosphate and nitrate/nitrite (measured together) for samples taken across the global oceans and at various depths. We also used the Tara Oceans dataset alongside the PISCES2 model (*30*, *31*) to estimate iron concentrations at each of these sites. Our statistical model yielded a list of genes with significant negative correlations (defined as a correlation coefficient cutoff of less than −0.7) to each of these nutrients (**Supplementary Table 2**) to ensure that only genes with a strong signal were included in our analysis.

Our resultant gene set (**Supplementary Table 2**) included a total of 24 genes with a strong negative correlation to either nitrate/nitrite, phosphate, or iron, meaning those genes are in high abundance when a given nutrient is present at low concentrations. Our analyses identified a much higher number of COGs with strong correlations to phosphate (16) compared to iron (3) and nitrate/nitrite (5), suggesting that modern marine microbial genomes are more strongly shaped by the availability of phosphate compared to iron or nitrate/nitrite. This list included known scavenging genes, such as *pst* genes encoding phosphate transporters that are particularly abundant in extremely low phosphate environments, and genes for synthesis and uptake of siderophores, which are compounds secreted by microorganisms that bind to iron to facilitate iron acquisition. The genes associated with low nitrate/nitrite concentrations were less obviously connected to nitrogen acquisition. For example, our list included genes such as a predicted mannosyl-3-phosphoglycerate phosphatase, which is a compatible solute that has been identified as an important protein stabilizer in both arid and hot environments (*32*, *33*). It also included a glycosidase, which is used in organic carbon processing and has been linked to an increase in organic matter content (*34*), though we were unable to find any studies linking these genes to nutrient limitation. One of the genes with a significant correlation to nitrate/nitrite also included *cbpA*, which is a DNA binding protein that is involved with microbial response to environmental stress: for example, it is involved in remodeling bacterial chromosomes in response to environmental stress in *E. coli* (*35*) and has been found to modulate a heat shock response in *Helicobacter pylori* (*36*). While none of these functions is directly related to nitrogen acquisition, the high abundance of these genes in low nitrogen environments suggests that they are part of a microbial physiological adaptation to persistent nitrogen limiting conditions.

The gene sets identified through these two approaches (a.k.a. the manual literature review and statistical analyses) had overlap, but were not identical to each other. Statistical analysis identified genes that are in high abundances when specific nutrients are low, and thus these are genes that are found in organisms adapted to persistently low nutrient levels, representing physiological adaptations to low nutrient regimes that could take a variety of forms. In contrast, genes identified through a literature review include those known to acquire specific nutrients.

This list includes genes that are found in a variety of nutrient regimes, but can be upregulated in response to a shift in nutrient availability. These would not be identified in our statistical analysis because our dataset was based on DNA rather than RNA sequence data. In this sense, genes identified based on metagenomic coverage in modern marine settings would include genes selected over longer periods of time, indicating adaptive evolution at the population level to sustained nutrient limitation. This is distinct from genes that are used for phenotypic plasticity, in which individuals up- or downregulate genes in response to transient changes in nutrient availability, which would likely be missed in the statistical analysis but would have been captured in the literature review.

It should also be noted that this analysis is based on data from marine settings, so would miss genes with significant correlations to nutrients that are exclusive to terrestrial or freshwater settings. Some COGs were eliminated from the final reconciliation analysis because they failed in our analysis pipeline either due to quality control or computational limitations: some COGs did not have sufficient data (i.e. fewer than 20 genes in the final alignment), some had poor alignments (i.e. passed our trimming and quality control steps with fewer than 100 amino acids), and some failed due to computational memory limitations during the reconciliation step (see **Supplementary Table 3** for a complete list of COGs, including those that failed in the pipeline).

### Identification of potential habitats for ancestral lineages

In order to determine the likely environments that these ancient lineages inhabited, we used reversible-jump Markov Chain Monte Carlo techniques to reconstruct the most probable habitat of ancestral lineages based on the habitats of their modern descendants (**Figure 1**). We restricted the analysis to include only three broad habitat types; shallow marine (i.e. above the photic zone), deep marine (i.e. below the photic zone), and terrestrial settings (including freshwater and human-associated habitats). We reconstructed the most probable habitat type for every internal node on our tree; **Figure 1** depicts the habitat reconstruction results for the major phyla. We used these results to ascertain the most likely habitat in which various gene events may have occurred to better understand which nutrients may have been limiting in which habitats. The reconstructed habitats were based on the modern habitats of thousands of archaea and bacteria, and were skewed towards terrestrial habitats as a result of modern sampling biases. Despite this, our reconstruction of ancestral habitats indicated that a substantial proportion of ancient lineages inhabited deep marine environments (**Figure 1**).

**Figure 1:**
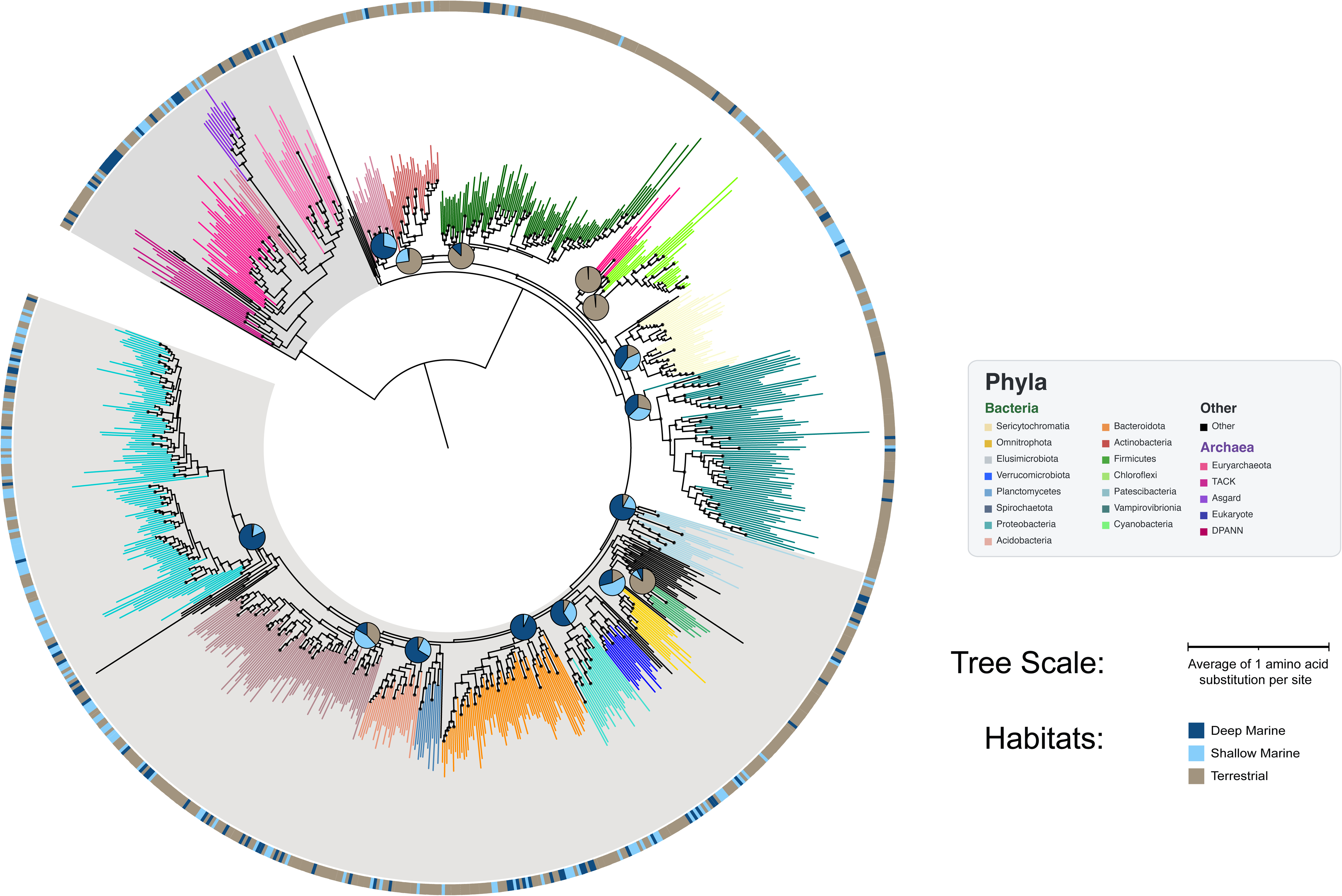
Tree of life (“species tree”) used for gene reconciliation analyses, showing reconstructed habitats for specific phyla over time. Phyla are color-coded and grouped to match to the larger phylum groups used for habitat reconstructions (see Methods). Outer ring shows habitat designations for all leaves of the species tree. Pie charts superimposed on internal nodes indicate the habitat reconstruction results of all organisms in the phylum (including additional genomes not represented in the species tree), with each wedge representing the probability that the internal node resided in a terrestrial (grey), shallow marine (light blue) or deep marine (dark blue) habitat. Habitat reconstruction results were based on much larger trees within each phylum (beyond those shown on the tree), so habitat reconstruction results were superimposed onto this tree for visualization purposes.

### Shifting nutrient availability in the early biosphere

Employing gene tree-species tree reconciliations, we reconstructed the complex evolutionary history of key genes associated with nutrient limitation, encompassing gene duplication, loss, horizontal gene transfer and speciation events. This analysis was complemented by reconstruction of the most probable habitat of ancestral lineages across deep marine, shallow marine, and terrestrial environments, thus contextualizing these evolutionary events within their likely ecological settings.

To isolate trends specific to our elements of interest (phosphorus, iron, nitrogen) and mitigate potential bioinformatics biases, we identified and reconciled all genes we identified to be associated with nutrient limitation (either through literature search or statistical analysis). Through this we were able to ascertain the timing of speciation events as well as gene duplications, losses, and horizontal gene transfer events at particular points in time. Our results are shown in **Figure 2**. **Figure 2A** illustrates gene events for genes identified through statistical analysis; **Figure 2B** illustrates speciations for genes identified through a literature search. Because horizontal gene transfer events, gene losses, and gene duplications occur on a branch between two nodes, we reported the midpoint date of the branch on which events occurred to summarize the broad trends of our results. We did not plot gene events that occurred on leaf branches, as these branches were extremely long and thus the timing was considerably more uncertain. We found that the relative patterns were consistent across all three clock models, but the timings for gene events reported by the CIR clock model were generally older than those identified by the LN and UGAM models. These figures below represent the timing for gene events according to the LN clock model because it was generally consistent with the UGAM model; **Supplementary Figure 2** shows the timing for gene events according to the CIR and UGAM clock models, and **Supplementary Figure 3** shows the timing for the full branch length along which these events could have occurred.

**Figure 2.**
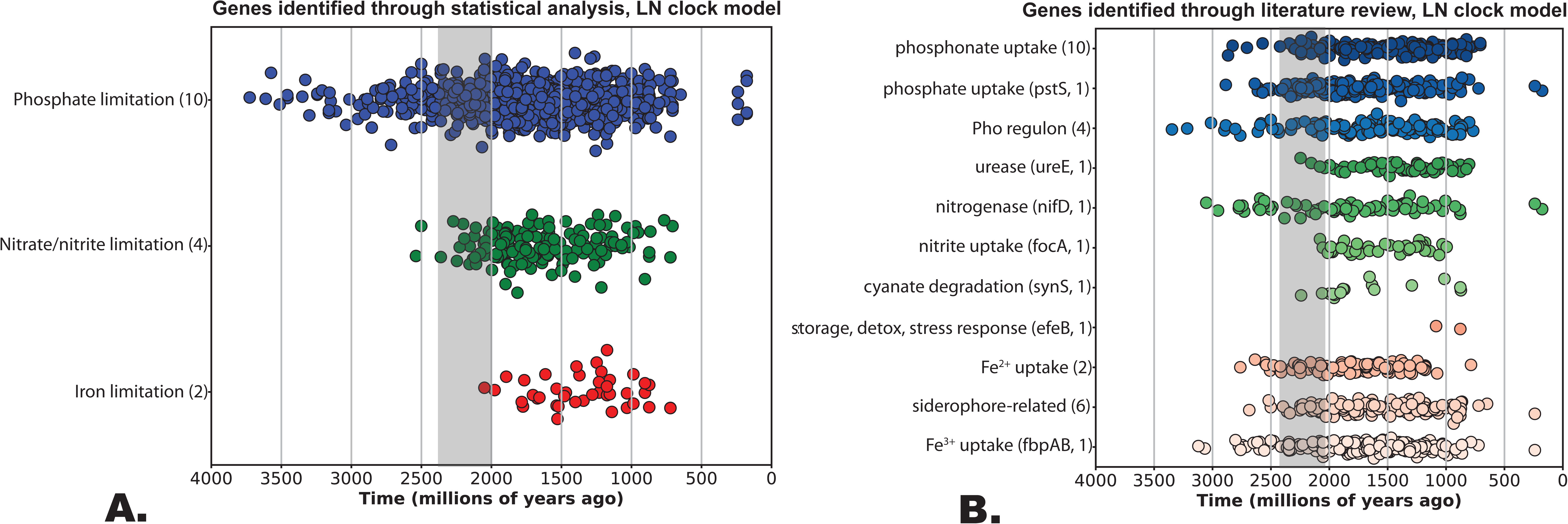
**Evolutionary history of gene events for genes associated with nutrient limitation relative to other genes**. Dots indicate gene events (speciations, horizontal gene transfer events, and duplications) for genes related to low concentrations of phosphate, nitrate/nitrite, and iron. Gene events (HGT, duplication) occur on tree branches; for this visualization, the date for a HGT or duplication gene event is reported based on the midpoint date of the node on which a gene event occurred. To see the full date range of the tree branches for these events, see **Supplementary** Figure 3. For this visualization, events occurring on terminal leaf nodes were removed because events occurring on long terminal branches artificially placed a cluster of events approximately 500 Mya. A) Genes identified through statistical analysis; only genes with a correlation slope of less than --0.7 indicating a large effect of nutrient concentration on gene abundance were plotted. B) Genes identified based on known function in the literature; genes are divided by functional category. A list and description of the genes included in this analysis are listed in **Supplementary Tables 1 and 2**. Further details are provided in the Methods. The Great Oxidation Event (GOE) is indicated with the grey bar. The events shown were calculated with the LN molecular clock model; other models are shown in the Supplementary Materials.

### Genes related to phosphate limitation are ancient

Our statistical analysis yielded a large number of COGs associated with phosphate limitation, suggesting that phosphate limitation leaves a more detectable imprint on modern microbial genomes compared to limitation of iron or nitrate/nitrate, based on the relative abundance of individual genes. Correspondingly, we found that these genes are relatively ancient, emerging as far back as the Archean era (**Figure 2**). Phosphorus scavenging genes identified via literature search showed similar trends, but closer analysis revealed that the most ancient genes associated with phosphorus limitation, dating back to at least the Archean era, are those associated with the phosphate (Pho) regulon (**Figure 2**). The Pho regulon is a set of genes associated with regulation and uptake of inorganic phosphate, and is found across a wide variety of bacteria (*37*). Genes in the Pho regulon are associated with the activation of extracellular enzymes capable of obtaining inorganic phosphorus from organic phosphorus compounds, phosphate transporters, and enzymes associated with P storage (*37*). These early-arising genes include *phoADX* (alkaline phosphatase) and *phoB* (phosphate stress response regulators). Previous work has indicated that secreted (as opposed to intracellular) PhoD alkaline phosphatases emerged around the Neoarchean or Paleoproterozoic (*38*), supporting these results. We also examined genes encompassing part of the phosphate-specific transport system (*pst*), including *pstABCS*. These ABC transporters use ATP to actively transport inorganic phosphate into the cell, are part of the Pho regulon and are known to be found in regions with particularly low phosphate concentrations (*26*). These genes emerged and radiated across the tree of life later than other genes in the Pho regulon, arising sometime between 3 Ga and 2.5 Ga according to the LN and UGAM clock models (**Figure 2, Supplementary Figures 2 and 3**). Their appearance later in the Archean relative to other P uptake genes is consistent with previous results (*26*), and may indicate that more severe P limitation emerged later in the Archean. Similarly, genes for uptake of phosphonate (an organic form of phosphorus) emerged later than genes related to the Pho regulon, at approximately the same time as (or a bit later than, depending on the clock model) genes related to phosphate uptake via *pst* (**Figure 2**). These results may imply that microorganisms initially dealt with P limitation by collecting organic P esters (or recycling organic P molecules inside their cells) via the *pho* genes, and only later employed ATP to actively pump Pi from the environment into their cells via the *pst* genes, perhaps in response to increasing P limitation.

**Figure 3:**
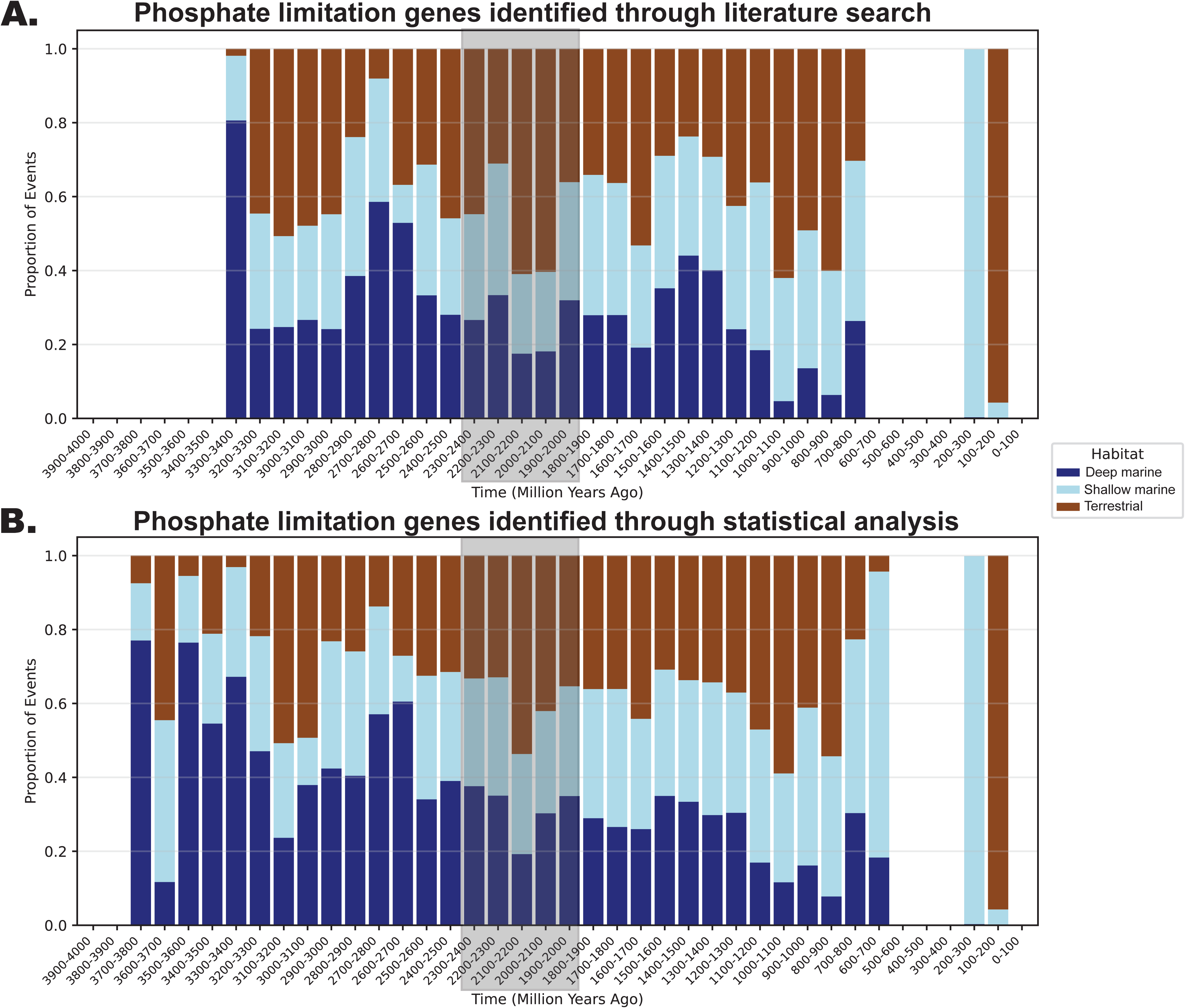
**Histograms of reconstructed habitats for gene events related to P limitation**. A., Putative habitats for gene events for P limitation genes identified through a literature search; B., Putative habitats for gene events for phosphate limitation genes identified through statistical analysis. GOE is indicated by a grey bar. Gene events are assigned a habitat based on the reconstructed habitat of the leftmost (earlier) node of the branch on which a gene event occurred, and the dates for the gene events are reported here based on the same node. The habitat for each node is assigned a posterior probability distribution; to create this figure, the posterior probability percentages of the predicted habitats for each gene event within a given time bin were added together and divided by the total number of gene events within the time bin to produce the percentages reported here. This graph depicts results from the left-hand node for the LN clock model. To see all histograms for all gene events for all three clock models and using dates from both the right- and left-hand nodes, see **Supplementary Figure 4.**

### A shift in nitrogen limitation through Earth history

Our statistical analysis yielded only 5 COGs with a significant negative correlation to nitrate/nitrite in modern oceans, and on geological timescales all of these COGs emerged well after those associated with phosphate limitation (**Figure 2**). Results from both the LN and UGAM clock models suggest that these genes emerged around the time of the Great Oxidation Event (GOE) in the Paleoproterozoic (ca. 2.4-2.1 Ga); the CIR clock placed their emergence earlier (**Supplementary Figures 2 and 3**). Genes identified via literature search are consistent with these results, suggesting that many genes associated with N limitation (nitrite uptake, urease, and cyanate degradation) did not emerge and begin to radiate across the tree of life until after the GOE. The genes identified in our analysis are not specific to the oxidized forms of nitrogen and instead reflect N limitation more broadly (from the literature review) or physiological responses to stress (from the statistical analysis), and thus do not reflect increased availability of oxidized nitrogen with the advent of the GOE. Instead, they suggest a broader limitation of nitrogen in the biosphere during the Proterozoic.

Importantly, this does not exclude the possibility that life acquired nitrogen through other means. For example, we were unable to include genes associated with ammonium uptake (particularly the Amt/Mep/Rh family of proteins) as well as many of the urease genes involved in urea uptake (particularly *ureABC*) in this study, because they failed in the reconciliation process due to computational constraints. However, we have recently used other methods to demonstrate that ammonia transporters emerged as far back as the last universal common ancestor (LUCA), ureases date back to the Archean, and nitrilases similarly emerged relatively early (*39*). Thus ammonium, urea, and nitriles could have been an important source of nitrogen to the early biosphere.

The advent of biological nitrogen fixation would have provided an important source of fixed nitrogen to the biosphere. We included reconciliation results for the nitrogenase subunit *nifD* in our analysis (**Figure 2**), which suggests that these genes emerged in the Mesoarchean (approximately 3 Ga, according to all three clock models), which is consistent with previous results (*40*). A more recent analysis found that these genes date back as far as LUCA; however, the various nitrogenase subunits required for nitrogen fixation *(nifD, nifK,* and *nifH*) may not have been united in the same organism until approximately 2.88-1.87 Ga, depending on the clock model (*39*). It is conceivable that the unification of these genes increased N_2_ fixation efficiency, perhaps in response to higher nitrogen demand. Importantly, those dates for the unification of *nif* genes roughly align with the timing for the emergence of other N limitation genes included in our analysis, consistent with increasing N-limitation at that time. Furthermore, this finding implies that N may not have been limiting during most of the Archean. If so, it would support the idea that some form of nitrogen, such as ammonium, cyanide or urea derived from biological or abiotic sources, was sufficiently bioavailable until at least the Neoarchean, while other nutrients, in particular phosphorus, were more limiting (*39*).

This rise in global N limitation during the Proterozoic could be linked to a nitrogen crisis triggered by the rise of microbial denitrification (*41*). The microbial process of denitrification, a microbial metabolism that converts dissolved nitrate into N_2_ gas, results in a loss of fixed nitrogen from the biosphere. We have previously dated the rise of denitrification to approximately 2.5 Ga (*40*), which would have deprived the biosphere of a vital nutrient in the form of nitrate. The genes that we identified through our literature search are known to be linked to N limitation, and the genes we identified via the statistical analysis likely represent physiological adaptations to persistently low nitrate. It is therefore striking that all of these genes seem to have emerged at approximately the same time, supporting the notion of a nitrogen crisis resulting from the rise of denitrification (*41*).

### Iron became more limiting after the Great Oxidation Event

Our statistical analysis yielded only 2 COGs that were significantly negatively correlated with iron concentrations according to the PISCES model, based on the Tara Oceans metagenomic data. These two COGs included genes related to iron transport and siderophores (**Supplementary Table 2**), and they did not emerge until well after genes associated with phosphate and nitrate/nitrite limitation in our analysis, after the GOE. Previous evolutionary studies on iron transport in Cyanobacteria are consistent with these results: transporters for Fe^2+^ and Fe^3+^ (FeoB, FutB and cFTR1) only stem back to in the Proterozoic (*42*). It has previously been hypothesized that as Earth became progressively more oxygenated, gigatons of soluble ferrous iron were removed from habitats due to precipitation of iron oxides (*43*). This would effectively decrease iron availability to the biosphere, thus increasing selection pressure for the acquisition of iron scavenging genes. As a consequence, the relative number of gene events for genes related to iron limitation increased relative to genes related to N or P limitation in the aftermath of the GOE. This is also consistent with mineralogical studies indicating millimolar iron levels in the Archean ocean (*44*) that likely decreased over time with decreasing hydrothermal inputs (*45*). In addition, DOC levels and microbial sulfate reduction likely expanded with atmospheric oxygenation and the growing biosphere (*1*, *4*), leading to Fe-scavenging along productive marine margins (*46*). Overall, there would thus have been increasing selection pressure for the emergence of iron acquisition strategies in the Proterozoic.

However, the genes that we identified through the literature tell a different story. We examined the evolutionary history of genes related to ferrous (Fe^2+^) and ferric (Fe^3+^) iron acquisition that had been identified in previous studies. Of particular interest were siderophores, which are high-affinity chelating molecules produced by a wide range of microbes to bind iron in the environment (*47*). Generally, siderophores are water-soluble molecules that are excreted into the environment, bind Fe^3+^, and then are taken back into the cell via specialized transporters.

Intracellular iron deficiency can trigger the expression of genes for siderophore biosynthesis and excretion, followed by uptake of iron-bound siderophores to replenish cellular iron stores (*47*, *48*). Intriguingly, our results indicate that genes associated with siderophore synthesis and uptake emerged early in the evolution of the biosphere, predating the Great Oxidation Event in all three of our clock models (arising approximately 2.7 Ga according to the LN and UGAM models, and as far back as ca. 3.0 Ga according to the CIR clock model) (**Figure 2; Supplementary Figures 2 and 3**). Similarly, other genes related to both Fe^2+^ and Fe^3+^ uptake appear to have emerged early, dating back to the Archean for all 3 clock models (ranging from ca. 2.7 Ga for Fe^2+^ and ca 3.2 Ga for Fe^3+^ transporters according to the LN clock; ca. 2.7 Ga for both Fe^2+^ and Fe^3+^ transporters according to the UGAM clock, and ca. 3.3 Ga for both Fe^2+^ and Fe^3+^ transporters for the CIR clock) (**Figure 2; Supplementary Figures 2 and 3**). However, these genes are associated with Fe uptake in general, and are not necessarily associated with Fe limitation.

The discrepancy between the timing for genes associated with low iron levels in modern oceans and those identified as iron scavenging genes based on a literature review may be due in part to the nature of the genes identified by each analysis. The list of genes with a significant negative correlation to iron concentrations included only one COG related to siderophores; the other three COGs were related to Fe^2+^ transport and hemin transport. Hemin (the oxidized form of heme) is a commonly occurring iron-containing organic molecule that can be released into the environment through organic matter degradation. Our results indicate that the abundance of siderophore-encoding genes generally does not correlate with iron concentrations in modern marine environments, suggesting that the ability to synthesize siderophores for iron acquisition is found even in organisms inhabiting iron-replete environments, and is likely upregulated in response to low iron concentrations. As a result, these results could suggest that selection favored the maintenance of iron-acquisition genes like siderophores early in Earth’s evolution because early siderophore-producing organisms inhabited environments with fluctuating iron concentrations. This finding is potentially consistent with the idea that Fe^2+^ was a limiting substrate for photoferrotrophs in the Archean ocean. Photoferrotrophs would have inhabited the photic zone, distal to deep-sea hydrothermal Fe sources, and they may have been among the dominant primary producers at that time (*14*) are often invoked as key agents in the production of iron ore deposits that sustain the modern economy (*49*). Hence their overall Fe demand would have been high, providing them with a competitive advantage if they were able to generate siderophores to overcome Fe limitation.

### Tracing nutrient limitation to specific habitat type

Because nutrient limitation on Earth today generally varies depending on location, we attempted to reconstruct the likely habitats of ancestral nodes using reversible-jump Markov Chain Monte Carlo simulation in order to identify the likely habitats in which these gene events occurred. This analysis would enable tracking of nutrient limitation with greater spatial resolution through time. However, our reconstruction of putative habitats for these gene events did not reveal markedly distinct trends for each of the gene categories we analyzed. While we urge caution in overinterpreting the reconstructed habitat results, the most interesting and noticeable trend we observed was a distinct decline in the proportion of gene events occurring in deep-sea habitats for genes related to P limitation (**Figure 3**; see **Supplementary Figure 4** for histograms of all gene categories and all three clock models). This was consistent for the sets of genes identified through both statistical analysis and literature review. In the modern ocean, P concentrations generally increase with depth (*50*), and our results may indicate that P was relatively more limiting in the deep ocean on the early Earth than it is today. This is perhaps consistent with the proposition that phosphate was scavenged by coprecipitation with Fe^2+^ minerals (vivanite, greenalite, green rust) in the deep ocean (*12*, *51*, *52*), while UV radiation in the surface ocean may have facilitated phosphate recycling (*53*). However, additional modeling and geochemical work is necessary to further support these results.

An additional noteworthy feature is that terrestrial habitats extend back to the beginning of the record in our datasets, although some geological models argue for much smaller continental land masses in the early Archean (*54*). In the absence of continents, the biosphere should have been almost entirely shifted into the marine realm. Furthermore, intense UV radiation on land prior to the establishment of the ozone shield during the GOE should have rendered terrestrial habitats less attractive to a thriving biosphere. Instead, our results tentatively suggest that terrestrial habitats date back to the early stages of life on Earth. Either small land masses were particularly attractive for microbial life, for example due to high nutrient supply, or land masses were more abundant than thought.

## Conclusions

On the modern Earth, the nutrient most responsible for limiting growth depends on the relationship between nutrient demand and nutrient availability. For example, in the modern ocean, upwelling regions are largely Fe-limited, nitrogen is limiting in subtropical gyres and in the Arctic Ocean in the summertime, and both nutrients can be co-limiting in intermediate regions (*55–57*). Phosphate is generally found in higher concentrations in the Pacific compared to the Atlantic Ocean, suggesting that N is more limiting in the Pacific compared to the Atlantic, where P is more limiting (*18*, *19*). Some organisms have different nutrient requirements compared to others; for example, diatoms have a high requirement of Si to construct their frustules. Our results should thus be interpreted in light of the complex and dynamic nature of nutrient limitation on the modern Earth: while nutrient limitation undoubtedly varied across habitat and by lineage, we have revealed patterns in global-scale nutrient limitation over Earth history. Our results suggest that natural selection has favored the maintenance of P uptake mechanisms since the Archean, whereas genes associated with the persistent limitation of N and Fe do not appear to have emerged until later in Earth history.

Nutrient limitation is inherently linked to biological elemental stoichiometry: in other words, nutrients become limiting for life when the relative abundance of that nutrient in the environment is low relative to the biological requirements for that nutrient (*8*, *10*, *58*). Thus, our results do not necessarily suggest that nitrate or iron became less abundant over time; instead, our results suggest that nitrate and iron became more limiting over time on a global scale relative to other nutrients required by life.

In this study, we identified genes that act as “sensors” for nutrient limitation using two distinct approaches: first, through identifying genes known to be associated with nutrient limitation based on the scientific literature; and second, a more agnostic approach employing a statistical model to identify genes linked to nutrient limitation by analyzing the relationship between gene abundances and nutrient concentrations in modern oceans. By exploring the evolutionary history of these gene paleosensors, we can infer which nutrients were likely the most limiting to the biosphere at various stages of Earth history. Our results suggest that P was a limiting nutrient for the biosphere much earlier than N and Fe, with persistent Fe limitation emerging only after the Great Oxidation Event.

Our results have important implications for understanding the scope of the biosphere and primary productivity as life and Earth co-evolved. Fluctuations in nutrient availability likely influenced the size of the biosphere in Earth’s oceans and on land, impacting biogeochemical cycles and major evolutionary transitions. Although further research is needed to compare these genomic results with modeling and geochemical records, our study provides a foundation for comparing and contextualizing other methods used to understand the evolution of Earth and its biosphere over time.

## Methods

### Identification of COGs associated with nutrient limitation using a statistical model

To identify clusters of orthologous genes (COGs) that correlate with nutrient limitation in the modern ocean, we examined the Ocean Microbial Reference Catalog v2 (OM-RGC.v2) from the *Tara* Oceans Project (*28*, *29*). The OM-RGC.v2 includes relative gene abundances of all COGs (*n* = 4,787) in 139 *Tara* Oceans metagenomic samples, along with metadata information including phosphate, oxygen, and nitrate/nitrite concentrations. (Nitrate/nitrite values were reported together for OM-RGC v2.) Iron concentrations for *Tara* Oceans samples were not available and were thus estimated using the PISCES2 model (*30*, *31*) based on iron concentration model predictions for *Tara* Oceans sampling locations as described in Table S1 of (*59*). Iron concentrations were predicted for surface and the deep chlorophyll maximum (DCM) only; iron concentrations for samples from the mesopelagic zone were not available under the PISCES2 model. All other metadata for *Tara* Oceans samples were directly obtained from (*29*).

Estimation of correlations between COGs and metadata information was performed using regression models. Compound poisson linear models were fitted in bulk using the MaAsLin2 software package (*60*) (v. 1.18.0). Separate models were fit for each COG to analyze the effect of metadata variables on individual COG abundances. While the main focus was to investigate correlation with nutrient abundance, environmental metadata was included in the model to control for as many potential confounding effects as the data allowed. The following predictors were included in the final model (based on variables available from the Tara Oceans dataset): the size fraction at which the sample was taken, mean temperature, depth, salinity, mean oxygen concentration, PO_4_ concentration, NO_2_ + NO_3_ concentration, iron concentration, and absolute latitude. Of these, the following predictors were log-transformed to allow greater model fit: depth, PO_4_ concentration, NO_2_ + NO_3_ concentration. To the same end, the iron concentration was transformed by taking the square root, and the absolute value of the latitude was taken. Otherwise, no transformations or normalization was performed. No abundance cutoff was applied, but COGs present in less than one-third of the Tara Oceans samples were discarded in order to ensure that the COGs identified by the statistical model were meaningful.

COGs were considered to be correlated with a given nutrient if the corresponding regression coefficient was found to be significant at a 0.05 significance level following a Benjamini-Hochberg correction procedure (*61*). In addition, correlations were considered strong if the sample Spearman correlation coefficient (*r)* between the abundances of the COG and the nutrient was less than −0.6, indicating that gene abundances were high when nutrient abundances were low. All statistical analyses were performed in R (*62*). A summary of the methods used to identify target genes and identify gene events is summarized in **Supplementary Figure 1.**

For the analysis distinguishing genes associated with ferric and ferrous iron, all COGs that were correlated with iron concentrations in our analysis were manually separated into COGs that bind to Fe^3+^ or Fe^2+^ based on Protein Data Bank (RCSB.org) (*63*) annotations and the primary literature. COGs that did not have a clear association with either Fe^3+^ or Fe^2+^ were not included for that analysis.

The code to perform this analysis is available at https://figshare.com/projects/_b_Time-resolved_phylogenomics_analysis_reveals_patterns_in_biosphere_nutrient_limitation_through_ Earth_history_b_/268163 (see maaslin_modelling.R). A full list of all the results from our statistical analysis of COG abundance with metadata collected by the Tara Oceans project is provided in this FigShare repository as well.

### Construction of a species tree and time-calibrated chronogram

In order to determine the evolutionary history of each COG of interest identified above, we first had to create a tree of life that is representative of the diversity of life on Earth, and anchor this tree in time to create a molecular clock. For this analysis, we used the phylogenetic tree and molecular clock previously described in (*26*). Briefly, the tree includes 865 different genomes chosen to represent the full diversity of bacteria and archaea. We used sixteen ribosomal genes that have been used previously to reconstruct the evolutionary history of life (*64*, *65*) to create the alignment that served as the basis for the final tree, as implemented in GToTree (*66*). The tree was created using maximum likelihood methodology implemented in IQ-Tree 2.2.0.3 (*67*). A relaxed Bayesian molecular clock was implemented in PhyloBayes 4.1 (*68*, *69*) with eight calibration points and a uniform root prior between 4.4 and 3.5 Ga. For further details regarding tree construction and molecular clock calibration, please see (*26*). Results shown in the main text are based on the LN model; results based on UGAM and CIR are shown in the Supplementary Materials.

### Identification of COGs within genomes

To reconstruct the evolutionary history of each COG, we had to identify those COGs within each of the genomes making up our species tree. To do this, the COG2020 database (*27*)], including the amino acid sequences of genes representing each COG, was downloaded from https://www.ncbi.nlm.nih.gov/research/cog-project/. DIAMOND (v0.9.22) (*70*) was used to align ORFs from the 865 genomes in our species tree to sequences in the COG database with a minimum 30% identity, maximum e-value of 10^-5^, minimum 70% subject and query alignment. If an ORF mapped to multiple sequences in the COG database, the match with the lowest e-value was chosen.

### Construction of gene trees for each COG

For each COG accession, we aligned the ORFs under this COG accession using MUSCLE v5.1 (*71*). From the multiple sequence alignment output, we first removed columns with >15% gaps across all sequences using TrimAL v1.4 (*72*) (using the command trimal -gt 0.85). Then we removed individual sequences that had greater than 20% gaps in the sequence after alignment.

For all COGs with ≥ 20 sequences remaining after alignment trimming, a gene tree was constructed for each COG accession number using IQ-TREE version 2.0.3 (*67*) using ultrafast bootstraps and the default UFBoot convergence criterion. COGs that failed to reach convergence after 3000 bootstraps were not analyzed. The model of evolution was determined by the Model Selection tool within IQ-TREE (*73*) with the flags *-m MFP, -mrate E,I,G,I+G,R and -madd C10,C20,C30,C40,C50,C60,EX2,EX3,EHO,UL2,UL3,EX_EHO,LG4M,LG4X,CF4,LG+C10,LG +C20,LG+C30,LG+C40,LG+C50,LG+C60* to include all complex mixture models of protein evolution in model selection.

### Reconciliation of gene trees with species trees

To identify gene duplications, losses, transfers, and speciation events, we reconciled gene trees with species trees using ecceTERA v1.2.5 (*74*). We used the default settings implemented in ecceTERA with the command amalgamate=true to consider uncertainty in the gene trees, and with transfers from the dead turned on. Reconciliation analyses were performed on fully dated species trees and full sets of ultrafast bootstraps for all gene trees. We calculated the mean date for each hgt and duplication event based on the midpoints of the two 95% confidence intervals that defined the nodes of the branch on which the event occurred. We used scripts provided in (*75*) to parse recPhyloXML files. Note that for all figures visualizing gene events, we did not include events occurring on leaf nodes. This is because leaf nodes were exceptionally long and many events occurred on leaf nodes, thus skewing the data visualizations by concentrating many events within the past 100 million years, a result that stemmed from the structure of the phylogenetic tree.

### Habitat assignment for ancestral lineages on the the tree of life

For this analysis, we used Bayesian trait prediction to reconstruct the likely habitats of ancestral lineages in the tree based on the known habitats of lineages included in our species tree. For further detail regarding how this analysis was conducted, please see the Supplementary Methods.

## Teaser

Phosphorus has been limiting since early in Earth history, while nitrogen and iron became limiting on a global scale after the Great Oxidation Event.

## Acknowledgements

This work was performed by the Virtual Planetary Laboratory team, a member of the NASA Nexus for Exoplanet System Science, funded by NASA Astrobiology Program Grant No. 80NSSC18K0829. Support was also provided by a Scialog program sponsored jointly by the Research Corporation for Science Advancement (RCSA) and the Heising-Simons Foundation and includes a grant (no. 28109) to Carleton College by RCSA. We also acknowledge funding from a NERC Frontiers grant (NE/V010824/1) to EES.

## Author contributions

Conceptualization: REA, EES

Investigation and analysis: ZN, TO, JZ, AG, WP, AK, BL, AL, JB, EES, REA

Visualization: ZN, REA, JB, TO, AG, WP, AK, BL

Supervision: REA, EES Writing: REA, EES, ZN, JB, TO

## Data and materials availability

All datasets, tables, and scripts associated with this study have been deposited in FigShare at https://figshare.com/projects/_b_Time-resolved_phylogenomics_analysis_reveals_patterns_in_biosphere_nutrient_limitation_through_ Earth_history_b_/268163.

## References

1. T. W. Lyons, C. J. Tino, G. P. Fournier, R. E. Anderson, W. D. Leavitt, K. O. Konhauser, E. E. Stüeken, Co-evolution of early Earth environments and microbial life. Nature Reviews Microbiology 22, 572–586 (2024).

2. N. J. Planavsky, S. A. Crowe, M. Fakhraee, B. Beaty, C. T. Reinhard, B. J. W. Mills, C. Holstege, K. O. Konhauser, Evolution of the structure and impact of Earth’s biosphere. Nat Rev Earth Environ 2, 123–139 (2021).

3. J.-W. Yang, M. Brandon, A. Landais, S. Duchamp-Alphonse, T. Blunier, F. Prié, T. Extier, Global biosphere primary productivity changes during the past eight glacial cycles. Science 375, 1145–1151 (2022).

4. P. W. Crockford, Y. M. B. On, L. M. Ward, R. Milo, I. Halevy, The geologic history of primary productivity. Current Biology 33, 4741–4750.e5 (2023).

5. J. Kallmeyer, R. Pockalny, R. R. Adhikari, D. C. Smith, S. D’Hondt, Global distribution of microbial abundance and biomass in subseafloor sediment. Proceedings of the National Academy of Sciences of the United States of America 109, 16213–6 (2012).

6. F. I. Woodward, Global primary production. Current Biology 17, R269–R273 (2007).

7. T. M. Hoehler, D. J. Mankel, P. R. Girguis, T. M. McCollom, N. Y. Kiang, B. B. Jørgensen, The metabolic rate of the biosphere and its components. Proceedings of the National Academy of Sciences 120, e2303764120 (2023).

8. A. Redfield, On the proportions of organic derivations in sea water and their relation to the composition of plankton.(1934) In James Johnstone Memorial Volume. University Press of Liverpool, Liverpool, 177–192 (1934).

9. T. Tyrrell, The relative influences of nitrogen and phosphorus on oceanic primary production. Nature 400, 525–531 (1999).

10. N. J. Planavsky, The elements of marine life. Nature Geoscience 7, 855–856 (2014).

11. J. J. Elser, M. E. S. Bracken, E. E. Cleland, D. S. Gruner, W. S. Harpole, H. Hillebrand, J. T. Ngai, E. W. Seabloom, J. B. Shurin, J. E. Smith, Global analysis of nitrogen and phosphorus limitation of primary producers in freshwater, marine and terrestrial ecosystems. Ecology Letters 10, 1135–1142 (2007).

12. C. T. Reinhard, N. J. Planavsky, B. C. Gill, K. Ozaki, L. J. Robbins, T. W. Lyons, W. W. Fischer, C. Wang, D. B. Cole, K. O. Konhauser, Evolution of the global phosphorus cycle. Nature 541, 386–389 (2017).

13. C. R. Walton, S. Ewens, J. D. Coates, R. E. Blake, N. J. Planavsky, C. Reinhard, P. Ju, J. Hao, M. A. Pasek, Phosphorus availability on the early Earth and the impacts of life. Nat. Geosci. 16, 399–409 (2023).

14. L. M. Ward, B. Rasmussen, W. W. Fischer, Primary Productivity Was Limited by Electron Donors Prior to the Advent of Oxygenic Photosynthesis. Journal of Geophysical Research: Biogeosciences 124, 211–226 (2019).

15. C. M. Moore, M. M. Mills, K. R. Arrigo, I. Berman-Frank, L. Bopp, P. W. Boyd, E. D. Galbraith, R. J. Geider, C. Guieu, S. L. Jaccard, T. D. Jickells, J. La Roche, T. M. Lenton, N. M. Mahowald, E. Marañón, I. Marinov, J. K. Moore, T. Nakatsuka, A. Oschlies, M. A. Saito, T. F. Thingstad, A. Tsuda, O. Ulloa, Processes and patterns of oceanic nutrient limitation. Nature Geosci 6, 701–710 (2013).

16. T. A. Laakso, D. P. Schrag, Limitations on Limitation. Global Biogeochemical Cycles 32, 486–496 (2018).

17. S. S. Merchant, J. D. Helmann, Elemental Economy: Microbial Strategies for Optimizing Growth in the Face of Nutrient Limitation. Advances in Microbial Physiology 60, 91–210 (2012).

18. A. C. Martiny, Y. Huang, W. Li, Occurrence of phosphate acquisition genes in Prochlorococcus cells from different ocean regions. Environmental Microbiology 11, 1340–1347 (2009).

19. L. J. Ustick, A. A. Larkin, C. A. Garcia, N. S. Garcia, M. L. Brock, J. A. Lee, N. A. Wiseman, J. Keith Moore, A. C. Martiny, J. K. Moore, A. C. Martiny, Metagenomic analysis reveals global-scale patterns of ocean nutrient limitation. Science 372, 287–291 (2021).

20. S. Lockwood, C. Greening, F. Baltar, S. E. Morales, Global and seasonal variation of marine phosphonate metabolism. ISME J 16, 2198–2212 (2022).

21. B. A. S. Van Mooy, G. Rocap, H. F. Fredricks, C. T. Evans, A. H. Devol, Sulfolipids dramatically decrease phosphorus demand by picocyanobacteria in oligotrophic marine environments. Proceedings of the National Academy of Sciences 103, 8607–8612 (2006).

22. G. Sharpe, L. Zhao, M. G. Meyer, W. Gong, S. M. Burns, A. Tagliabue, K. N. Buck, A. E. Santoro, J. R. Graff, A. Marchetti, S. Gifford, Synechococcus nitrogen gene loss in iron-limited ocean regions. ISME Communications 3, 107 (2023).

23. C. A. Garcia, G. I. Hagstrom, A. A. Larkin, L. J. Ustick, S. A. Levin, M. W. Lomas, A. C. Martiny, Linking regional shifts in microbial genome adaptation with surface ocean biogeochemistry. Philosophical Transactions of the Royal Society B: Biological Sciences 375, 20190254 (2020).

24. M. Fakhraee, L. G. Tarhan, C. T. Reinhard, N. J. Planavsky, Constraining the elemental stoichiometry of early marine life: Geology, v. 51. Geological Society of America | GEOLOGY 51, 1043–1047 (2023).

25. J. E. Horne, C. Goldblatt, EONS: A New Biogeochemical Model of Earth’s Oxygen, Carbon, Phosphorus, and Nitrogen Systems From the Archean to the Present. Geochemistry, Geophysics, Geosystems 25, e2023GC011252 (2024).

26. J. S. Boden, J. Zhong, R. E. Anderson, E. E. Stüeken, Timing the evolution of phosphorus-cycling enzymes through geological time using phylogenomics. Nat Commun 15, 3703 (2024).

27. M. Y. Galperin, Y. I. Wolf, K. S. Makarova, R. V. Alvarez, D. Landsman, E. V. Koonin, COG database update: Focus on microbial diversity, model organisms, and widespread pathogens. Nucleic Acids Research 49, D274–D281 (2021).

28. S. Sunagawa, L. P. Coelho, S. Chaffron, J. R. Kultima, K. Labadie, G. Salazar, B. Djahanschiri, G. Zeller, D. R. Mende, A. Alberti, F. M. Cornejo-castillo, P. I. Costea, C. Cruaud, F. Ovidio, S. Engelen, I. Ferrera, J. M. Gasol, L. Guidi, F. Hildebrand, F. Kokoszka, C. Lepoivre, Structure and function of the global ocean microbiome. 348, 1–9 (2015).

29. G. Salazar, L. Paoli, A. Alberti, J. Huerta-Cepas, H. J. Ruscheweyh, M. Cuenca, C. M. Field, L. P. Coelho, C. Cruaud, S. Engelen, A. C. Gregory, K. Labadie, C. Marec, E. Pelletier, M. Royo-Llonch, S. Roux, P. Sánchez, H. Uehara, A. A. Zayed, G. Zeller, M. Carmichael, C. Dimier, J. Ferland, S. Kandels, M. Picheral, S. Pisarev, J. Poulain, S. G. Acinas, M. Babin, P. Bork, E. Boss, C. Bowler, G. Cochrane, C. de Vargas, M. Follows, G. Gorsky, N. Grimsley, L. Guidi, P. Hingamp, D. Iudicone, O. Jaillon, S. Kandels-Lewis, L. Karp-Boss, E. Karsenti, F. Not, H. Ogata, S. Pesant, N. Poulton, J. Raes, C. Sardet, S. Speich, L. Stemmann, M. B. Sullivan, S. Sunagawa, P. Wincker, Gene Expression Changes and Community Turnover Differentially Shape the Global Ocean Metatranscriptome. Cell 179, 1068–1083.e21 (2019).

30. O. Aumont, C. Ethé, A. Tagliabue, L. Bopp, M. Gehlen, PISCES-v2: an ocean biogeochemical model for carbon and ecosystem studies. Geoscientific Model Development 8, 2465–2513 (2015).

31. A. Tagliabue, O. Aumont, R. DeAth, J. P. Dunne, S. Dutkiewicz, E. Galbraith, K. Misumi, J. K. Moore, A. Ridgwell, E. Sherman, C. Stock, M. Vichi, C. Völker, A. Yool, How well do global ocean biogeochemistry models simulate dissolved iron distributions? Global Biogeochemical Cycles 30, 149–174 (2016).

32. M. S. da Costa, H. Santos, E. A. Galinski, “An overview of the role and diversity of compatible solutes in Bacteria and Archaea” in Biotechnology of Extremophiles, G. Antranikian, Ed. (Springer, Berlin, Heidelberg, 1998; 10.1007/BFb0102291), pp. 117–153.

33. C. Li, H. Liao, L. Xu, C. Wang, M. Yao, J. Wang, X. Li, Comparative genomics reveals the adaptation of ammonia-oxidising Thaumarchaeota to arid soils. Environmental Microbiology 26, e16601 (2024).

34. B. H. Hill, C. M. Elonen, T. M. Jicha, A. M. Cotter, A. S. Trebitz, N. P. Danz, Sediment microbial enzyme activity as an indicator of nutrient limitation in Great Lakes coastal wetlands. Freshwater Biology 51, 1670–1683 (2006).

35. K. Chintakayala, S. S. Singh, A. E. Rossiter, R. Shahapure, R. T. Dame, D. C. Grainger, E. coli Fis Protein Insulates the cbpA Gene from Uncontrolled Transcription. PLOS Genetics 9, e1003152 (2013).

36. D. Roncarati, A. Danielli, V. Scarlato, CbpA Acts as a Modulator of HspR Repressor DNA Binding Activity in Helicobacter pylori. Journal of Bacteriology 193, 5629–5636 (2011).

37. F. Santos-Beneit, The Pho regulon: a huge regulatory network in bacteria. Front. Microbiol. 6 (2015).

38. A. Sanger, A. D. Steen, J. S. Boden, S. Cellier-Goetghebeur, M. Bayder, E. P. Mueller, G. P. Halverson, N. B. Cowan, R. E. Anderson, J. N. Pelletier, E. E. Stüeken, M. Fakhraee, K. O. Konhauser, N. Mahmoudi, Microbial exoenzymes catalyzed the transition to an oxygenated Earth. bioRxiv [Preprint] (2025). 10.1101/2025.09.11.675588.

39. J. Boden, Z. Ni, R. E. Anderson, E. E. Stuueken, Abiotic Sources of Fixed Nitrogen Sustained Early Ecosystems for Several Hundred MIllion Years After the Origin of Life. Research Square [Preprint] (2025). 10.21203/rs.3.rs-7759286/v1.

40. C. Parsons, E. E. Stüeken, C. J. Rosen, K. Mateos, R. E. R. E. Anderson, E. E. Stueeken, C. J. Rosen, K. Mateos, R. E. R. E. Anderson, Radiation of nitrogen-metabolizing enzymes across the tree of life tracks environmental transitions in Earth history. Geobiology 19, 18–34 (2021).

41. K. Fennel, M. Follows, P. G. Falkowski, The Co-Evolution of the Nitrogen, Carbon and Oxygen Cycles in the Proterozoic Ocean. American Journal of Science 305, 526–545 (2005).

42. T. C. Enzingmüller-Bleyl, J. S. Boden, A. J. Herrmann, K. W. Ebel, P. Sánchez-Baracaldo, N. Frankenberg-Dinkel, M. M. Gehringer, On the trail of iron uptake in ancestral Cyanobacteria on early Earth. Geobiology 20, 776–789 (2022).

43. J. Wade, D. J. Byrne, C. J. Ballentine, H. Drakesmith, Temporal variation of planetary iron as a driver of evolution. Proceedings of the National Academy of Sciences of the United States of America 118, e2109865118 (2021).

44. N. J. Tosca, S. Guggenheim, P. K. Pufahl, An authigenic origin for Precambrian greenalite: Implications for iron formation and the chemistry of ancient seawater. GSA Bulletin 128, 511–530 (2016).

45. E. E. S. S. Viehmann, S. Viehmann, J. Krayer, S. Weyer, E. E. Stu00fceken, S. V. Hohl, N. Tepe, Y. Lin, D. Kraemer, D. M. Ernst, M. V. Kranendonk, Europium traces the impact of high temperature hydrothermal systems on the early oceans. Geochemical Perspectives Letters 34, 57–61 (2025).

46. S. W. Poulton, P. W. Fralick, D. E. Canfield, Spatial variability in oceanic redox structure 1.8 billion years ago. Nature Geosci 3, 486–490 (2010).

47. I. J. Schalk, Bacterial siderophores: diversity, uptake pathways and applications. Nat Rev Microbiol 23, 24–40 (2025).

48. R. Saha, N. Saha, R. S. Donofrio, L. L. Bestervelt, Microbial siderophores: a mini review. Journal of Basic Microbiology 53, 303–317 (2013).

49. K. O. Konhauser, T. Hamade, R. Raiswell, R. C. Morris, F. G. Ferris, G. Southam, D. E. Canfield, Could bacteria have formed the Precambrian banded iron formations? Geology 30, 1079–1082 (2002).

50. L. A. Bristow, W. Mohr, S. Ahmerkamp, M. M. M. Kuypers, Nutrients that limit growth in the ocean. Current Biology 27, R474–R478 (2017).

51. M. P. Brady, R. Tostevin, N. J. Tosca, Marine phosphate availability and the chemical origins of life on Earth. Nat Commun 13, 5162 (2022).

52. I. Sugiyama, I. Halevy, Interaction of metal oxyions and phosphate with carbonate green rust: Insights into Earth’s modern and ancient environments. Geochimica et Cosmochimica Acta 397, 96–112 (2025).

53. O. Farr, J. Hao, W. Liu, N. Fehon, J. R. Reinfelder, N. Yee, P. G. Falkowski, Archean phosphorus recycling facilitated by ultraviolet radiation. Proceedings of the National Academy of Sciences 120, e2307524120 (2023).

54. C. Hawkesworth, P. A. Cawood, B. Dhuime, T. Kemp, Tectonic processes and the evolution of the continental crust. Journal of the Geological Society 181, jgs2024-027 (2024).

55. J. H. Martin, Glacial-interglacial CO2 change: The Iron Hypothesis. Paleoceanography 5, 1–13 (1990).

56. M. J. Behrenfeld, Z. S. Kolber, Widespread Iron Limitation of Phytoplankton in the South Pacific Ocean. Science 283, 840–843 (1999).

57. T. J. Browning, C. M. Moore, Global analysis of ocean phytoplankton nutrient limitation reveals high prevalence of co-limitation. Nat Commun 14, 5014 (2023).

58. M. Fakhraee, L. G. Tarhan, C. T. Reinhard, N. J. Planavsky, Constraining the elemental stoichiometry of early marine life. Geology 51, 1043–1047 (2023).

59. L. Caputi, Q. Carradec, D. Eveillard, A. Kirilovsky, E. Pelletier, J. J. Pierella Karlusich, F. Rocha Jimenez Vieira, E. Villar, S. Chaffron, S. Malviya, E. Scalco, S. G. Acinas, A. Alberti, J.-M. Aury, A.-S. Benoiston, A. Bertrand, T. Biard, L. Bittner, M. Boccara, J. R. Brum, C. Brunet, G. Busseni, A. Carratalà, H. Claustre, L. P. Coelho, S. Colin, S. D’Aniello, C. Da Silva, M. Del Core, H. Doré, S. Gasparini, F. Kokoszka, J.-L. Jamet, C. Lejeusne, C. Lepoivre, M. Lescot, G. Lima-Mendez, F. Lombard, J. Lukeš, N. Maillet, M.- A. Madoui, E. Martinez, M. G. Mazzocchi, M. B. Néou, J. Paz-Yepes, J. Poulain, S. Ramondenc, J.-B. Romagnan, S. Roux, D. Salvagio Manta, R. Sanges, S. Speich, M. Sprovieri, S. Sunagawa, V. Taillandier, A. Tanaka, L. Tirichine, C. Trottier, J. Uitz, A. Veluchamy, J. Veselá, F. Vincent, S. Yau, S. Kandels-Lewis, S. Searson, C. Dimier, M. Picheral, T. O. Coordinators, P. Bork, E. Boss, C. de Vargas, M. J. Follows, N. Grimsley, L. Guidi, P. Hingamp, E. Karsenti, P. Sordino, L. Stemmann, M. B. Sullivan, A. Tagliabue, A. Zingone, L. Garczarek, F. d’Ortenzio, P. Testor, F. Not, M. R. d’Alcalà, P. Wincker, C. Bowler, D. Iudicone, Community-Level Responses to Iron Availability in Open Ocean Plankton Ecosystems. Global Biogeochemical Cycles 33, 391–419 (2019).

60. H. Mallick, A. Rahnavard, L. J. McIver, S. Ma, Y. Zhang, L. H. Nguyen, T. L. Tickle, G. Weingart, B. Ren, E. H. Schwager, S. Chatterjee, K. N. Thompson, J. E. Wilkinson, A. Subramanian, Y. Lu, L. Waldron, J. N. Paulson, E. A. Franzosa, H. C. Bravo, C. Huttenhower, Multivariable association discovery in population-scale meta-omics studies. PLOS Computational Biology 17, e1009442 (2021).

61. Y. Benjamini, Y. Hochberg, Controlling the False Discovery Rate: A Practical and Powerful Approach to Multiple Testing. Journal of the Royal Statistical Society: Series B (Methodological) 57, 289–300 (1995).

62. R Core Team, R: A Language and Environment for Statistical Computing (R Foundation for Statistical Computing, Vienna, Austria, 2025; https://www.R-project.org/).

63. H. M. Berman, J. Westbrook, Z. Feng, G. Gilliland, T. N. Bhat, H. Weissig, I. N. Shindyalov, P. E. Bourne, The Protein Data Bank. Nucleic Acids Res 28, 235–242 (2000).

64. L. A. Hug, B. J. Baker, K. Anantharaman, C. T. Brown, A. J. Probst, C. J. Castelle, C. N. Butterfield, A. W. Hernsdorf, Y. Amano, I. Kotaro, Y. Suzuki, N. Dudek, D. A. Relman, K. M. Finstad, R. Amundson, B. C. Thomas, J. F. Banfield, A new view of the tree and life’s diversity. Nature Microbiology 1, 16048 (2016).

65. C. J. Castelle, J. F. Banfield, Major New Microbial Groups Expand Diversity and Alter our Understanding of the Tree of Life. Cell 172, 1181–1197 (2018).

66. M. D. Lee, GToTree: a user-friendly workflow for phylogenomics. Bioinformatics 35, 4162–4164 (2019).

67. B. Q. Minh, H. A. Schmidt, O. Chernomor, D. Schrempf, M. D. Woodhams, A. von Haeseler, R. Lanfear, IQ-TREE 2: New Models and Efficient Methods for Phylogenetic Inference in the Genomic Era. Mol Biol Evol 37, 1530–1534 (2020).

68. N. Lartillot, T. Lepage, S. Blanquart, PhyloBayes 3: A Bayesian software package for phylogenetic reconstruction and molecular dating. Bioinformatics 25, 2286–2288 (2009).

69. N. Lartillot, PhyloBayes : Bayesian Phylogenetics Using Site-heterogeneous Models. 0–16 (2020).

70. B. Buchfink, C. Xie, D. H. Huson, Fast and sensitive protein alignment using DIAMOND. Nat Methods 12, 59–60 (2015).

71. R. C. Edgar, Muscle5: High-accuracy alignment ensembles enable unbiased assessments of sequence homology and phylogeny. Nat Commun 13, 6968 (2022).

72. S. Capella-Gutiérrez, J. M. Silla-Martínez, T. Gabaldón, trimAl: a tool for automated alignment trimming in large-scale phylogenetic analyses. Bioinformatics 25, 1972–1973 (2009).

73. S. Kalyaanamoorthy, B. Q. Minh, T. K. F. Wong, A. von Haeseler, L. S. Jermiin, ModelFinder: fast model selection for accurate phylogenetic estimates. Nature Methods 14, 587–589 (2017).

74. E. Jacox, C. Chauve, G. J. Szöllősi, Y. Ponty, C. Scornavacca, ecceTERA: comprehensive gene tree-species tree reconciliation using parsimony. Bioinformatics 32, 2056–2058 (2016).

75. W. Duchemin, G. Gence, A. M. A. Chifolleau, L. Arvestad, M. S. Bansal, V. Berry, B. Boussau, F. Chevenet, N. Comte, A. A. Davín, C. Dessimoz, D. Dylus, D. Hasic, D. Mallo, R. Planel, D. Posada, C. Scornavacca, G. Szöllősi, L. Zhang, É. Tannier, V. Daubin, RecPhyloXML: A format for reconciled gene trees. Bioinformatics 34, 3646–3652 (2018).

