## Supplementary Materials for "Time-resolved phylogenomics analysis reveals patterns in biosphere nutrient limitation through Earth history"

**The supplementary materials include the following information:**
Supplementary Tables 1-3
Supplementary Figures 1-6
Supplementary Methods

Extended data, including datasets and scripts, can be found on FigShare at https://figshare.com/projects/_b_Time-resolved_phylogenomics_analysis_reveals_patterns_in_biosphere_nutrient_limitation_through_Earth_history_b_/268163

| Gene Name | Limiting Nutrient | Function category | Ferrous or Ferric iron | COG ID | Function | Source |
|---|---|---|---|---|---|---|
| FbpAB | Iron | non-siderophore Fe3 transporter | Ferric | COG1178 | ABC-type $Fe^{3+}$ transport system, permease component | Moraes et al., 2021 |
| FepA | Iron | siderophore-related | Ferric | COG4771 | Outer membrane receptor for ferrienterochelin and colicins | Malmstrom et al., 2013 |
| EntC | Iron | siderophore-related | Ferric | COG1169 | Isochorismate synthase EntC | Lamb et al., 2016 |
| EntD | Iron | siderophore-related | Ferric | COG2977 | 4'-phosphopantetheinyl transferase EntD (siderophore biosynthesis) | Raymond et al., 2003; Noinaj et al., 2011 |
| EntF | Iron | siderophore-related | Ferric | COG3319 | Non-ribosomal peptide synthetase component F/Thioesterase domain of type I polyketide synthase or non-ribosomal peptide synthetase | Raymond et al., 2003; Noinaj et al., 2011 |
| FecA | Iron | siderophore-related | Ferric | COG4772 | Encodes a TBDT for ferric citrate; part of the fecABCDE operon. Outer membrane receptor | Noinaj et al., 2011 |
| FhuE | Iron | siderophore-related | Ferric | COG4773 | Outer membrane receptor for ferric coprogen and ferric-rhodotorulic acid. | Noinaj et al., 2011 |
| FTR1 | Iron | direct Fe2 uptake | Ferrous | COG0672 | High-affinity Fe2+/Pb2+ permease | Lau et al., 2016 |
| FeoA | Iron | direct Fe2 uptake | Ferrous | COG1918 | Fe2+ transport system protein FeoA | Shin et al., 2012 |

| | | | | | | |
|---|---|---|---|---|---|---|
| EfeB | Iron | storage, detox, and stress response | Ferrous | COG2837 | Periplasmic deferrochelatase/peroxidase EfeB | Cao et al., 2017 |
| cynS | Nitrogen | cyanate degradation | | COG1513 | Encodes cyanate lyase. | Flores et al., 2001 |
| focA | Nitrogen | nitrite uptake | | COG2116 | formate transporter/nitrite transporter | Shi et al., 2009 |
| nifDK | Nitrogen | nitrogenase | | COG2710 | nitrogenase nifD | |
| ureE | Nitrogen | urease | | COG2371 | Encodes urease accessory protein UreE. | Flores et al., 2001 |
| phoA | Phosphorus | Pho regulon | | COG1785 | Alkaline phosphatase (PhoA) | Chisholm et al., 2006; Chisholm et al., 2010; Hudek et al., 2016 |
| phoB | Phosphorus | Pho regulon | | COG1785 | Phosphate stress response regulator | Chisholm et al., 2006; Hudek et al., 2016 |
| phoD | Phosphorus | Pho regulon | | COG3540 | Alkaline phosphatase (PhoD) | Santos Beneit, 2015 |
| phoX | Phosphorus | Pho regulon | | COG3211 | Alkaline phosphatase (PhoX) | Balan et al., 2017 |
| pstS | Phosphorus | phosphate uptake | | COG0226 | ABC-type phosphate transport system, periplasmic component | Chisholm et al., 2006 |
| phnC | Phosphorus | phosphonate uptake | | COG3638 | ABC-type phosphonate transport system, periplasmic component | Chisholm et al., 2006; Stasi et al., 2019 |
| phnE1 | Phosphorus | phosphonate uptake | | COG3639 | ABC-type phosphonate transport system, periplasmic component | Chisholm et al., 2006; Stasi et al., 2019 |
| phnE2 | Phosphorus | phosphonate uptake | | COG3639 | ABC-type phosphonate transport system, periplasmic component | Chisholm et al., 2006; Stasi et al., 2019 |
| phnG | Phosphorus | phosphonate uptake | | COG3624 | ABC-type phosphonate transport system, periplasmic component | Brodersen et al., 2015 |
| phnH | Phosphorus | phosphonate uptake | | COG3625 | ABC-type phosphonate transport system, periplasmic component | Jia et al., 2008; Brodersen et al., 2015 |
| phnI | Phosphorus | phosphonate uptake | | COG3626 | ABC-type phosphonate transport system, periplasmic component | Brodersen et al., 2015 |
| phnL | Phosphorus | phosphonate uptake | | COG4778 | ABC-type phosphonate transport system, periplasmic component | Brodersen et al., 2015 |
| phnM | Phosphorus | phosphonate uptake | | COG3454 | ABC-type phosphonate transport system, periplasmic component | Boden et al., 2024 |

| | | | | | | |
|---|---|---|---|---|---|---|
| phnN | Phosphorus | phosphonate uptake | | COG3709 | ABC-type phosphonate transport system, periplasmic component | Boden et al., 2024 |
| phnJ | Phosphorus | phosphonate uptake | | COG3627 | ABC-type phosphonate transport system, periplasmic component | Brodersen et al., 2015; Boden et al., 2024 |

**Supplementary Table 1**. All COGs identified as being associated with limitation of phosphorus, nitroge, and iron based on literature review.

| COG Number | Correlated variable | Correlation coefficient | COG letter | COG category | COG annotation |
|---|---|---|---|---|---|
| COG3470 | iron | -0.7304334 | MI | Cell wall/membrane/envelope biogenesis; Lipid transport and metabolism; | Uncharacterized protein probably involved in high-affinity Fe2+ transport |
| COG4558 | iron | -0.7207392 | P | Inorganic ion transport and metabolism; | ABC-type hemin transport system, periplasmic component |
| COG2308 | nitrate/nitrite | -0.781165049 | S | Function unknown; | Uncharacterized conserved protein, circularly permuted ATPgrasp superfamily |
| COG3769 | nitrate/nitrite | -0.776617953 | G | Carbohydrate transport and metabolism; | Predicted mannosyl-3-phosphoglycerate phosphatase, HAD superfamily |
| COG2307 | nitrate/nitrite | -0.75182854 | S | Function unknown; | Uncharacterized conserved protein, Alpha-E superfamily |
| COG0366 | nitrate/nitrite | -0.702948524 | G | Carbohydrate transport and metabolism; | Glycosidase |
| COG0226 | phosphate | -0.893171234 | P | Inorganic ion transport and metabolism; | ABC-type phosphate transport system, periplasmic component |
| COG3484 | phosphate | -0.835445245 | O | Post-translational modification, protein turnover, and chaperones; | Predicted proteasome-type protease |
| COG0855 | phosphate | -0.785420155 | P | Inorganic ion transport and metabolism; | Polyphosphate kinase |

| | | | | | |
|---|---|---|---|---|---|
| COG3769 | phosphate | -0.742298417 | G | Carbohydrate transport and metabolism; | Predicted mannosyl-3-phosphoglycerate phosphatase, HAD superfamily |
| COG3651 | phosphate | -0.740546174 | S | Function unknown; | Uncharacterized conserved protein, DUF2237 family |
| COG1828 | phosphate | -0.720258254 | F | Nucleotide transport and metabolism; | Phosphoribosylformylglycinamidine (FGAM) synthase, PurS component |
| COG0400 | phosphate | -0.717585841 | R | General function prediction only; | Predicted esterase |
| COG5470 | phosphate | -0.71092123 | S | Function unknown; | Uncharacterized conserved protein, DUF1330 family |
| COG0248 | phosphate | -0.707519501 | FTP | Nucleotide transport and metabolism; Signal transduction mechanisms; Inorganic ion transport and metabolism; | Exopolyphosphatase/pppGpp-phosphohydrolase |
| COG0717 | phosphate | -0.702099164 | F | Nucleotide transport and metabolism; | Deoxycytidine triphosphate deaminase |

**Supplementary Table 2**. List of all COGs identified having statistically significant, strong correlations (correlation coefficient greater than 0.7) in abundance with phosphate, nitrate/nitrite, and iron according to the Tara Oceans dataset.

| **COG ID** | **Function** | **Gene Name** | **Limiting Nutrient** | **Identified through statistical analysis or literature review?** | **Status** |
|---|---|---|---|---|---|
| COG1178 | ABC-type $Fe^{3+}$ transport system, permease component | FbpAB | Iron | Literature review | Included in final list |
| COG4771 | Outer membrane receptor for ferrienterochelin and colicins | FepA | Iron | Literature review | Included in final list |
| COG0811 | Biopolymer transport protein ExbB/TolQ | TolQ | Iron | Literature review | Failed in ecceTERA step due to memory limitations |

| | | | | | |
|---|---|---|---|---|---|
| COG0848 | Biopolymer transport protein ExbD - associated with tonB | ExbD | Iron | Literature review | Failed in ecceTERA step due to memory limitations |
| COG1169 | Isochorismate synthase EntC | EntC | Iron | Literature review | Included in final list |
| COG2977 | 4'-phosphopantetheinyl transferase EntD (siderophore biosynthesis) | EntD | Iron | Literature review | Included in final list |
| COG3319 | Non-ribosomal peptide synthetase component E (peptide arylation enzyme) | EntF | Iron | Literature review | Included in final list |
| COG1120 | ABC-type cobalamin/Fe3+-siderophores transport system, ATPase component | FepC | Iron | Literature review | Failed in ecceTERA step due to memory limitations |
| COG0609 | ABC-type Fe3+-siderophore transport system, permease component | FepD | Iron | Literature review | Failed in ecceTERA step due to memory limitations |
| COG4772 | Encodes a TBDT for ferric citrate; part of the fecABCDE operon. Outer membrane receptor | FecA | Iron | Literature review | Included in final list |
| COG4773 | Outer membrane receptor for ferric coprogen and ferric-rhodotorulic acid. | FhuE | Iron | Literature review | Included in final list |
| COG0672 | High-affinity Fe2+/Pb2+ permease | FTR1 | Iron | Literature review | Included in final list |
| COG1918 | Fe2+ transport system protein FeoA | FeoA | Iron | Literature review | Included in final list |
| COG0735 | $Fe^{2+}$ or $Zn^{2+}$ uptake regulation protein | Fur | Iron | Literature review | Failed in ecceTERA step due to memory limitations |
| COG4592 | ABC-type Fe2+-enterobactin transport system, periplasmic component | FepB | Iron | Literature review | Removed in pipeline due to quality checks: fewer than 20 ORFs in final alignment. |
| COG2837 | Periplasmic deferrochelatase/peroxidase EfeB | EfeB | Iron | Literature review | Included in final list |

| | | | | | |
|---|---|---|---|---|---|
| COG1513 | Encodes cyanate lyase. | cynS | Nitrogen | Literature review | Included in final list |
| COG2116 | formate transporter/nitrite transporter | focA | Nitrogen | Literature review | Included in final list |
| COG2710 | nitrogenase nifD | nifDK | Nitrogen | Literature review | Included in final list |
| COG2371 | Encodes urease accessory protein UreE. | ureE | Nitrogen | Literature review | Included in final list |
| COG1785 | Alkaline phosphatase (PhoA) | phoA | Phosphoru s | Literature review | Included in final list |
| COG1785 | Phosphate stress response regulator | phoB | Phosphoru s | Literature review | Included in final list |
| COG3540 | Alkaline phosphatase (PhoD) | phoD | Phosphoru s | Literature review | Included in final list |
| COG3211 | Alkaline phosphatase (PhoX) | phoX | Phosphoru s | Literature review | Included in final list |
| COG0581 | ABC-type phosphate transport system, permease component | pstA | Phosphoru s | Literature review | Failed in ecceTERA step due to memory limitations |
| COG1117 | ABC-type phosphate transport system, ATPase component | pstB | Phosphoru s | Literature review | Failed in ecceTERA step due to memory limitations |
| COG0573 | ABC-type phosphate transport system, permease component | pstC | Phosphoru s | Literature review | Failed in ecceTERA step due to memory limitations |
| COG0226 | ABC-type phosphate transport system, periplasmic component | pstS | Phosphoru s | Literature review | Included in final list |
| COG3638 | ABC-type phosphonate transport system, periplasmic component | phnC | Phosphoru s | Literature review | Included in final list |
| COG3221 | ABC-type phosphonate transport system, periplasmic component | phnD | Phosphoru s | Literature review | Failed in ecceTERA step due to memory limitations |
| COG3639 | ABC-type phosphonate transport system, periplasmic component | phnE1 | Phosphoru s | Literature review | Included in final list |
| COG3639 | ABC-type phosphonate | phnE2 | Phosphoru s | Literature review | Included in final list |

| | | | | | |
|---|---|---|---|---|---|
| | transport system, periplasmic component | | | | |
| COG3624 | ABC-type phosphonate transport system, periplasmic component | phnG | Phosphoru s | Literature review | Included in final list |
| COG3625 | ABC-type phosphonate transport system, periplasmic component | phnH | Phosphoru s | Literature review | Included in final list |
| COG3626 | ABC-type phosphonate transport system, periplasmic component | phnI | Phosphoru s | Literature review | Included in final list |
| COG4778 | ABC-type phosphonate transport system, periplasmic component | phnL | Phosphoru s | Literature review | Included in final list |
| COG3454 | ABC-type phosphonate transport system, periplasmic component | phnM | Phosphoru s | Literature review | Included in final list |
| COG3709 | ABC-type phosphonate transport system, periplasmic component | phnN | Phosphoru s | Literature review | Removed in pipeline due to quality checks: fewer than 20 ORFs in final alignment. |
| COG4693 | Oxidoreductase (NAD-binding), involved in siderophore biosynthesis | PchG | iron | Literature review | Removed in pipeline due to quality checks: fewer than 20 ORFs in final alignment. |
| COG3627 | ABC-type phosphonate transport system, periplasmic component | phnJ | Phosphoru s | Literature review | Included in final list |
| COG0609 | ABC-type Fe3+-siderophore transport system, permease component | | iron | Statistical analysis | Failed in ecceTERA step due to memory limitations |

| | | | | | |
|---|---|---|---|---|---|
| COG3470 | Uncharacterized protein probably involved in high-affinity Fe2+ transport | | iron | Statistical analysis | Included in final list |
| COG4558 | ABC-type hemin transport system, periplasmic component | | iron | Statistical analysis | Included in final list |
| COG2308 | Uncharacterized conserved protein, circularly permuted ATPgrasp superfamily | | nitrate/nitrite | Statistical analysis | Included in final list |
| COG3769 | Predicted mannosyl-3-phosphoglycerate phosphatase, HAD superfamily | | nitrate/nitrite | Statistical analysis | Included in final list |
| COG2307 | Uncharacterized conserved protein, Alpha-E superfamily | | nitrate/nitrite | Statistical analysis | Included in final list |
| COG2214 | Curved DNA-binding protein CbpA, contains a DnaJ-like domain | | nitrate/nitrite | Statistical analysis | Removed in pipeline due to quality checks: fewer than 20 ORFs in final alignment. |
| COG0366 | Glycosidase | | nitrate/nitrite | Statistical analysis | Included in final list |
| COG0573 | ABC-type phosphate transport system, permease component | | phosphate | Statistical analysis | Failed in ecceTERA step due to memory limitations |
| COG0581 | ABC-type phosphate transport system, permease component | | phosphate | Statistical analysis | Failed in ecceTERA step due to memory limitations |
| COG1117 | ABC-type phosphate transport system, ATPase component | | phosphate | Statistical analysis | Failed in ecceTERA step due to memory limitations |
| COG0226 | ABC-type phosphate transport system, periplasmic component | | phosphate | Statistical analysis | Included in final list |

| | | | | | |
|---|---|---|---|---|---|
| COG3484 | Predicted proteasome-type protease | | phosphate | Statistical analysis | Included in final list |
| COG0855 | Polyphosphate kinase | | phosphate | Statistical analysis | Included in final list |
| COG0745 | DNA-binding response regulator, OmpR family, contains REC and winged-helix (wHTH) domain | | phosphate | Statistical analysis | Failed in ecceTERA step due to memory limitations |
| COG3769 | Predicted mannosyl-3-phosphoglycerate phosphatase, HAD superfamily | | phosphate | Statistical analysis | Included in final list |
| COG3651 | Uncharacterized conserved protein, DUF2237 family | | phosphate | Statistical analysis | Included in final list |
| COG1828 | Phosphoribosylformylglycinamidine (FGAM) synthase, PurS component | | phosphate | Statistical analysis | Included in final list |
| COG0400 | Predicted esterase | | phosphate | Statistical analysis | Included in final list |
| COG0413 | Ketopantoate hydroxymethyltransferase | | phosphate | Statistical analysis | Failed in ecceTERA step due to memory limitations |
| COG5470 | Uncharacterized conserved protein, DUF1330 family | | phosphate | Statistical analysis | Included in final list |
| COG0248 | Exopolyphosphatase/pppGpp-phosphohydrolase | | phosphate | Statistical analysis | Included in final list |
| COG1942 | Phenylpyruvate tautomerase PptA, 4-oxalocrotonate tautomerase family | | phosphate | Statistical analysis | Removed in pipeline due to quality checks: alignment less than 100 a.a.s |
| COG0717 | Deoxycytidine triphosphate deaminase | | phosphate | Statistical analysis | Included in final list |

**Supplementary Table 3**. All COGs included in our analysis, including those that failed in the pipeline due to quality control or computational limitations.

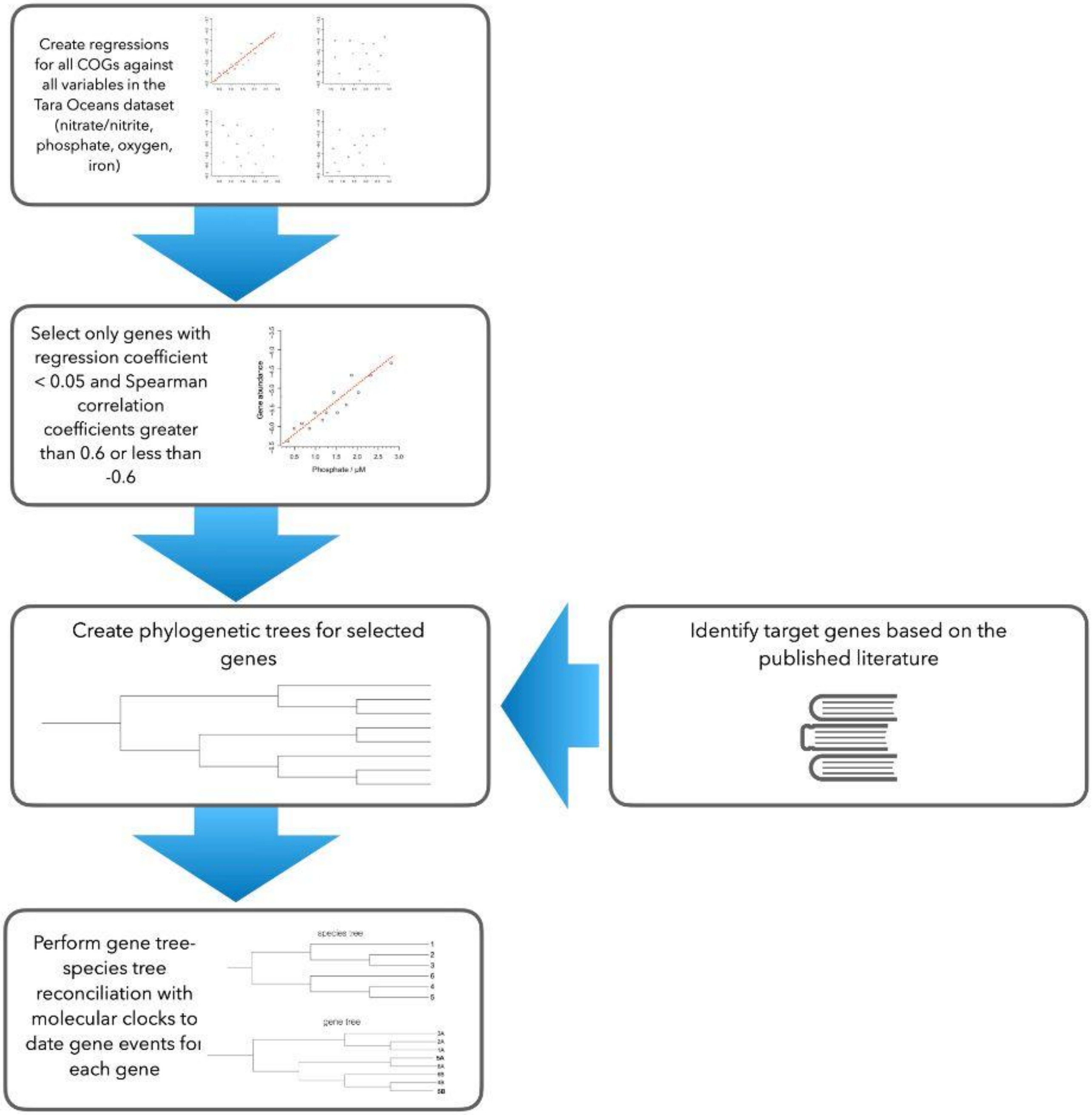


**Supplementary Figure 1**. Flow chart of methods used in this paper.

**Supplementary Figure 2**. **Evolutionary history of gene events for genes associated with nutrient limitation relative to other genes**. Dot plots show the number of gene events (speciations, horizontal gene transfer events, and duplications) for genes related to low concentrations of phosphate, nitrate/nitrite, and iron. The date for the gene event is reported based on the midpoint date of the node on which a gene event occurred. A) Genes identified through statistical analysis; only genes with a correlation slope of less than --0.7 were plotted. B) Genes identified through based on known function in the literature; genes are divided by functional category. Genes included in this analysis are listed in Supplementary Tables 1 and 2. Further details are provided in the Methods. The Great Oxidation Event (GOE) is indicated with the grey bar. The events shown were calculated with all three clock models.

**Supplementary Figure 3.** Diagrams indicating the full branch length along which gene horizontal gene transfer, duplication, and speciation events may have occurred. Genes identified through literature review are shown on the left; genes identified through statistical analysis are shown on the right. Gene events occurring on leaf nodes are faded in color. Gene event timings for all three molecular clock models (LN, CIR and UGAM) are depicted. Great Oxidation Event is depicted as a gray bar in all graphs.

**Supplementary Figure 4**: **Histograms of reconstructed habitats for gene events related to P limitation**. A., Putative habitats for gene events for all genes identified through a literature search; B., Putative habitats for gene events for all genes identified through statistical analysis. Gene events are assigned a habitat based on the reconstructed habitat of the leftmost (earlier) node of the branch on which a gene event occurred, and the dates for the gene events are reported here based on the same node. The habitat for each node is assigned a posterior probability distribution; to create this figure, the posterior probability percentages of the predicted habitats for each gene event within a given time bin were added together and divided by the total number of gene events within the time bin to produce the percentages reported here. This graph depicts results from both the left-hand and right-hand node for all three clock models.

### Supplementary Methods

#### *Habitat assignment for ancestral lineages on the tree of life*

##### Habitat identification for leaves in tree

Metadata and textual descriptions of habitat information were obtained by cross-referencing several databases using the GenBank accession numbers of the genomes used to construct the species tree. These databases include NCBI BioProject, BioSample, Genomes OnLine Database (GOLD) v.8: Public (*1*), Release 214 of GTDB metadata (*2*), Bergey's Manual of Systematic Bacteriology, and the published papers describing the original sample collection. For genomes from the Tara Oceans project, habitat information was assigned based on the sample site where the MAG was most abundant based on mapping of raw reads from to individual sample sites to the assembled contigs for each MAG (the original analysis for which these datasets were generated is described in (*3*). Based on the habitat information, we manually assigned each genome included in the tree to one of three broad habitat groups: deep marine, shallow marine, or terrestrial. Deep marine habitats are defined as aphotic zones of the ocean water column, including the mesopelagic layer, at depths greater than 100 m. Shallow marine habitats are defined as the marine euphotic zone, including the deep chlorophyll maximum layer, at depths less than 100 m. Terrestrial habitats are defined as non-marine environments, such as freshwater and sediment layers. Host-associated samples were assigned the same habitat as their host (e.g., bacterial genomes from human samples were categorized as "terrestrial"). The resulting dataset consists of habitat meta-information paired with a 3-class habitat label (see dataset posted on FigShare at https://figshare.com/projects/_b_Time-resolved_phylogenomics_analysis_reveals_patterns_in_biosphere_nutrient_limitation_through_Earth_history_b_/268163).

##### Habitat prediction for internal nodes in the tree

In order to predict the likely habitat for internal nodes of the species tree with greater confidence, we expanded the dataset to include 13,637 additional genomes so that more data could be used to predict the habitat for internal nodes in the 865-leaf species tree that we used for gene-species tree reconciliation. In order to assign a habitat to each of those additional genomes, we used the above 865 genome dataset as a training set to develop a model that predicts the habitat of the new expanded dataset, based on concatenated habitat information collected in the same way as the original, manually curated habitat dataset. We utilized the Term Frequency-Inverse Document Frequency (TF-IDF) method (*4*, *5*) for feature extraction to convert textual descriptions into numerical vectors. A total of 3,801 feature vectors were extracted to capture the textual habitat information. An ensemble method, the Extreme Gradient Boosting (XGBoost) Classifier algorithm (*6*), was used for habitat classification. The data was split into training, evaluation, and test sets with a ratio of 0.9:0.05:0.05. Hyperparameter tuning was performed using 5-fold stratified cross-validation. The best hyperparameters identified were max_depth=6, learning_rate=0.01, n_estimators=350, subsample=0.7, and colsample_bytree=0.7, which were used to train the final XGBoost classifier. The confusion matrix and the resulting Receiver Operating Characteristic (ROC) curves for the multi-class classification problem are shown in Supplementary Fig. 5, with an average area under the ROC curve (AUC) of 0.987. AUC quantifies the performance of a classifier by measuring its ability to distinguish between classes. The AUC ranges from 0.5 (random performance) to 1.0 (perfect discrimination), where higher values indicate better classification accuracy. The predicted habitats for each strain are available on FigShare at https://figshare.com/projects/_b_Time-resolved_phylogenomics_analysis_reveals_patterns_in_biosphere_nutrient_limitation_through_Earth_history_b_/268163.

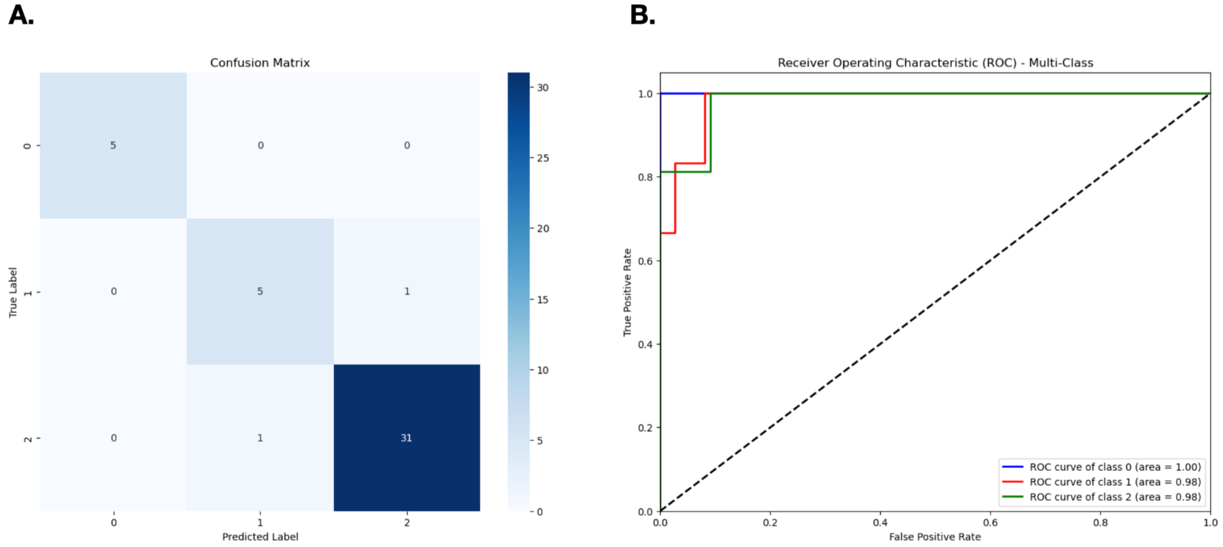


**Supplementary Fig. 5. Evaluation of habitat prediction performance.** A., Confusion matrix for three habitat classes (terrestrial, shallow marine, deep marine) showing 41/43 correct predictions (95.3% accuracy); B., One-vs-rest ROC curves show strong separability (AUC: terrestrial = 1.00; shallow marine = 0.98; deep marine = 0.98). The dashed diagonal indicates chance performance (true positive rate = false positive rate; AUC = 0.5).

#### Phylum-level ancestral habitat reconstruction

*Construction of phylum-level species tree*

Having classified the habitats of the additional genomes using the machine learning method described above, we used these additional genomes to better predict the habitat of the internal nodes on our species tree. The habitat reconstruction was separated for all major bacterial phyla to improve granularity and account for differences in evolutionary rates. To do this, we created new sub-trees for individual phyla with these additional genomes, predicted the habitats of internal nodes, then matched the internal nodes of the more detailed phyla trees with the original 865-leaf species tree we used for gene-species tree reconciliation.

To construct the individual phylum sub-trees, phyla forming monophyletic clades in the original 865-leaf tree of life were identified. For polyphyletic clades, phyla were assigned based on those contributing more than 10% of the total genomes in that clade. To enhance habitat diversity within each identified phylum group, additional genomes were included by sampling one genome from each GTDB representative genus under the phylum groups, resulting in a total of 13,637 additional genomes, divided into 16 phyla groups (annotated as Pseudonomadota, Chloroflexi, and so on). Homologs of 16 ribosomal proteins (L18, L3, L5, S8, L4, S3_C, L6, L2, L15e, S19, S17, L22, S10, L24, L16, and L14) previously used to infer the tree of life (*7–9*) were identified in each genome using HMMER3 v3.3.2 (hmmer.org); 95% of the genomes contained homologs of eight or more of these genes. Genomes with less than 50% of the markers were removed. The sequences were then aligned and trimmed to remove columns containing more than 15% gaps using MUSCLE v5.1 (*10*) and TrimAl v1.4.rev15 (*11*), all implemented in GToTree v1.6.34 (*12*). The resulting alignments were used to reconstruct a species tree using maximum likelihood as implemented in IQ-TREE v2.0.3 (*13*). Partitioned analyses were applied to allow each protein to evolve under appropriate substitution models. To determine these models, ModelFinder (*14*) was implemented with default options (no mixture models) and 'MERGE' to test whether merging individual partitions increased model fit. For all phylum-level species trees, the best model was found to be a single partition containing all proteins; therefore, the LG substitution rate matrix was applied to all 16 ribosomal protein alignments, with 10 categories of the FreeRate model to estimate substitution rates and their variations at different sites. To ensure consistency between the phyla-specific trees and the 865-leaf tree of life, we constrained the phylogenetic relationships among the original taxa from each phylum group to match their topology in the tree of life when inferring each expanded phylum-specific tree. An archaeal outgroup was included for rooting each phylum tree. The super5 algorithm in MUSCLE was used for trees consisting of over 1,000 genomes.

*Reconstruction of ancestral habitats*

BayesTraits (*15*) was used to reconstruct the habitat of all internal nodes in each phylum-specific tree. We used reverse-Jump MCMC with an exponential HyperPrior with a mean ranging from 0 to 46.43. The chains were run for 5 million iterations with sampling every 1,000 iterations. Samples from the last 500,000 iterations were used for the analysis. The MCMC log file was analyzed with the R package CODA (v0.19-4) (*16*). Heidelberger and Welch's diagnostic was performed to test whether the sampled values had reached a stationary distribution and the accuracy of mean estimation, with the Benjamini-Hochberg procedure to control the false discovery rate. For each internal node, we used the maximum a posteriori probability to assign the habitat. Full statistics of the MCMC chains are reported on FigShare at https://figshare.com/projects/_b_Time-resolved_phylogenomics_analysis_reveals_patterns_in_biosphere_nutrient_limitation_through_Earth_history_b_/268163.

This method of habitat reconstruction inferred putative habitats for each internal node of the species tree. Gene- and species-tree reconciliation identifies putative gene events on the internal branches of the tree. The habitat for individual gene events was depicted in figures to be the habitat for the left-hand node of the branch on which a gene event occurred.

*Ablation study for prior choice*

To test the effect of prior choice on habitat reconstruction results, a hyperparameter search was performed to select the priors for the Bayesian MCMC analysis. The search grid was constructed for the upper bound of the mean of the HyperPrior, with values incrementing at fixed intervals from 0 to 5, then at 1.25-fold intervals up to 100 for thoroughness. The resulting MCMC runs used the same iterations and were analyzed using the same metrics. To compare the reconstruction results, we generated the distribution of habitats for all nutrients examined, evolutionary event types, and habitats. Kullback-Leibler (KL) divergence was used to quantify the divergence between distributions, aggregated for all nutrients examined, evolutionary event types, and habitats to compare between two reconstructions (Supplementary Figure 6). Priors from clusters in the dendrograms resulted in qualitatively similar patterns of habitat evolution, indicating that the analysis was robust to the choice of priors.

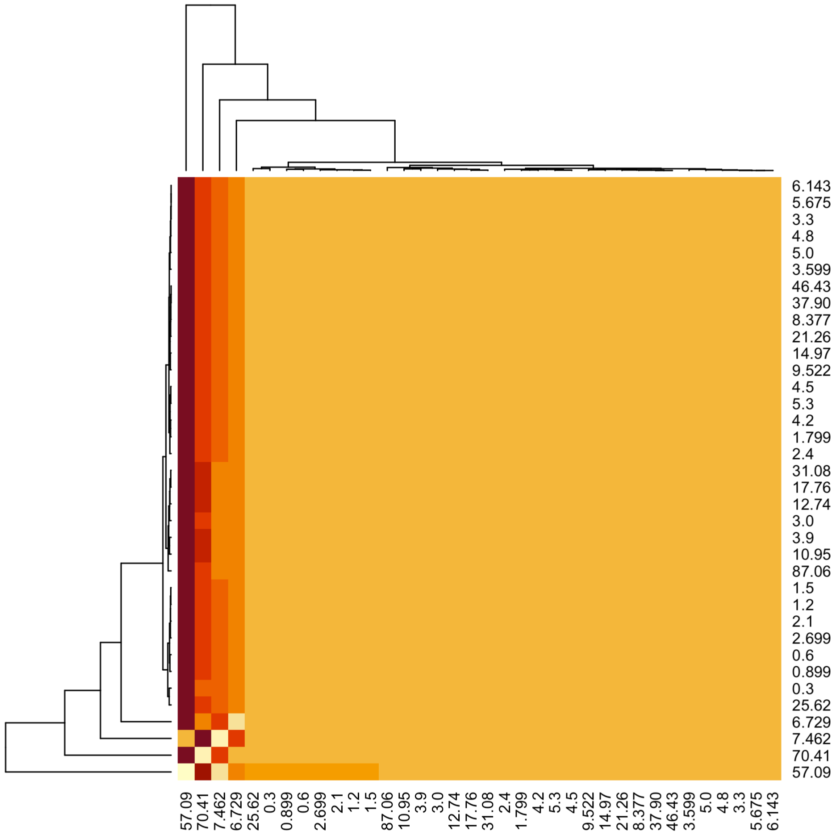


**Supplementary Figure 6. Clustered heatmap of KL divergence between reconstructions under different prior choice.** The plot shows pairwise KL divergence between results from ancestral habitat reconstructions using different priors (listed on row and column axes). Hierarchical clustering (dendrograms, left and top) groups priors based on the similarity of their results. Darker color indicates higher degree of dissimilarity. The dendrogram reveals distinct clusters of similar reconstructions.

#### Supplementary References


1. S. Mukherjee, D. Stamatis, C. T. Li, G. Ovchinnikova, M. Kandimalla, V. Handke, A. Reddy, N. Ivanova, T. Woyke, E. A. Eloe-Fardosh, I.-M. A. Chen, N. C. Kyrpides, T. B. K. Reddy, Genomes OnLine Database (GOLD) v.10: new features and updates. *Nucleic Acids Res* **53**, D989–D997 (2025).

2. D. H. Parks, M. Chuvochina, C. Rinke, A. J. Mussig, P.-A. Chaumeil, P. Hugenholtz, GTDB: an ongoing census of bacterial and archaeal diversity through a phylogenetically consistent, rank normalized and complete genome-based taxonomy. *Nucleic Acids Res* **50**, D785–D794 (2022).

3. J. Zhong, T. Osborn, T. Del Rosario Hernández, O. Kyrysyuk, B. J. Tully, R. E. Anderson, Increasing transposase abundance with ocean depth correlates with a particle-associated lifestyle. *mSystems* **9**, e00067-24 (2024).

4. L. Havrlant, V. Kreinovich, A simple probabilistic explanation of term frequency-inverse document frequency (tf-idf) heuristic (and variations motivated by this explanation). *International Journal of General Systems* **46**, 27–36 (2017).

5. K. S. Jones, A statistical interpretation of term specificity and its application in retrieval. *Journal of Documentation* **28** (1972).

6. T. Chen, C. Guestrin, "XGBoost: A Scalable Tree Boosting System" in *Proceedings of the 22nd ACM SIGKDD International Conference on Knowledge Discovery and Data Mining* (Association for Computing Machinery, New York, NY, USA, 2016; https://dl.acm.org/doi/10.1145/2939672.2939785)*KDD '16*, pp. 785–794.

7. L. A. Hug, B. J. Baker, K. Anantharaman, C. T. Brown, A. J. Probst, C. J. Castelle, C. N. Butterfield, A. W. Hernsdorf, Y. Amano, I. Kotaro, Y. Suzuki, N. Dudek, D. A. Relman, K. M. Finstad, R. Amundson, B. C. Thomas, J. F. Banfield, A new view of the tree and life's diversity. *Nature Microbiology* **1**, 16048 (2016).

8. K. Mateos, G. Chappell, A. Klos, B. Le, J. Boden, E. Stüeken, R. Anderson, The evolution and spread of sulfur cycling enzymes reflect the redox state of the early Earth. *SCIENCE ADVANCES*.

9. J. S. Boden, J. Zhong, R. E. Anderson, E. E. Stüeken, Timing the evolution of phosphorus-cycling enzymes through geological time using phylogenomics. *Nat Commun* **15**, 3703 (2024).

10. R. C. Edgar, MUSCLE: multiple sequence alignment with high accuracy and high throughput. *Nucleic Acids Research* **32**, 1792–1797 (2004).

11. S. Capella-Gutierrez, J. M. Silla-Martinez, T. Gabaldon, trimAl: a tool for automated alignment trimming in large-scale phylogenetic analyses. *Bioinformatics* **25**, 1972–1973 (2009).

12. M. D. Lee, GToTree: a user-friendly workflow for phylogenomics. *Bioinformatics* **35**, 4162–4164 (2019).

13. B. Q. Minh, H. A. Schmidt, O. Chernomor, D. Schrempf, M. D. Woodhams, A. von Haeseler, R. Lanfear, IQ-TREE 2: New Models and Efficient Methods for Phylogenetic Inference in the Genomic Era. *Mol Biol Evol* **37**, 1530–1534 (2020).

14. S. Kalyaanamoorthy, B. Q. Minh, T. K. F. Wong, A. von Haeseler, L. S. Jermiin, ModelFinder: fast model selection for accurate phylogenetic estimates. *Nature Methods* **14**, 587–589 (2017).

15. A. Meade, M. Pagel, "Ancestral State Reconstruction Using BayesTraits" in *Environmental Microbial Evolution: Methods and Protocols*, H. Luo, Ed. (Springer US, New York, NY, 2022; https://doi.org/10.1007/978-1-0716-2691-7_12), pp. 255–266.

16. M. Plummer, N. Best, K. Cowles, K. Vines, CODA: Convergence Diagnosis and Output Analysis for MCMC. *R News* **6**, 7–11 (2006).
